# Mixed origin of juvenile Atlantic cod (*Gadus morhua*) along the Swedish west coast

**DOI:** 10.1101/2022.06.03.494672

**Authors:** Simon Henriksson, Ricardo T. Pereyra, Marte Sodeland, Olga Ortega-Martinez, Halvor Knutsen, Håkan Wennhage, Carl André

## Abstract

Cryptic population structure in exploited fish species poses a major challenge for fisheries management. Atlantic cod (*Gadus morhua*) is a species in which the presence of sympatric ecotypes has been known for a long time, for instance off the coast of Northern Norway. More recently, two sympatric ecotypes of cod have also been documented in the Skagerrak and Kattegat; one ecotype is of an apparent offshore origin and undertakes spawning migrations to the North Sea, and the other is resident at the coast throughout its life. However, their relative contributions of juveniles to the Swedish west coast remain poorly understood. The lack of adult cod along the Skagerrak and Kattegat coasts in recent years has led to the hypothesis that the offshore ecotype is the main source of juveniles to the area, but recent studies have shown large proportions of coastal cod inside Norwegian Skagerrak fjords. In this study, juvenile cod were collected at a high spatial resolution along the Swedish west coast, and genetically assigned to each of the two ecotypes. The results reveal that there is a considerable proportion of juvenile coastal cod in the southern Kattegat, Öresund, and in inshore Swedish Skagerrak, but that the offshore ecotype dominates in offshore areas. Model selection suggests that differences in bottom depth, rather than distance from the open sea, may explain the heterogenous spatial distribution of the two ecotypes. In addition, the two ecotypes displayed differences at loci known to be associated with environmental adaptation, suggesting that their spatial distribution is maintained by natural selection in response to specific environmental conditions.

## Introduction

Overexploitation of fish stocks is cause for great concern for economic, socio-political, and ecological reasons, motivating a revision of management strategies (Pauly *et al*., 2002; FAO, 2020). Inaccurate delineation of management units, or “stocks”, has been suggested as one explanation for the failed management of many fish species (Reiss *et al*., 2009). Historical definitions of management units have often been based on legislative borders, despite knowledge that distribution ranges of marine populations may range from highly local to transboundary (Kerr *et al*., 2017). Management regimes that do not consider the population structure at the correct spatiotemporal scale may thus be suboptimal, potentially leading to underexploitation of productive populations and overexploitation of vulnerable populations (Kerr *et al*., 2017). Overexploitation may cause a loss of genetic diversity, reducing the species’ robustness and elevating the risk of extinction (Allendorf *et al*., 2022).

Populations with heritable differences in behaviour, morphology, physiology, or life history traits associated with adaptation to different environments are often referred to as “ecotypes” (Stronen *et al*., 2022), and have been described in several fish species (e.g. herring *[Clupea harengus],* Bekkevold *et al*., 2016; three-spined stickleback *[Gasterosteus aculeatus],* Hohenlohe *et al*., 2010; and Atlantic cod *[Gadus morhua],* Michalsen *et al*., 2014; Knutsen *et al*., 2018). One apparently common form of divergence in marine fishes is the evolution of offshore and coastal ecotypes, as described for polar cod *(Boreogadus saida,* Madsen *et al*., 2016), beaked redfish *(Sebastes mentella,* Cadrin *et al*., 2010), European seabass *(Dicentrarcus labrax,* Allegrucci *et al*., 1997), and European anchovy *(Engraulis encrasicolus*, Le Moan *et al*., 2016).

Atlantic cod has been a highly important commercial fish species in the North Atlantic since at least the 16^th^ century (COSEWIC, 2010). In recent times, stocks on both sides of the Atlantic have been severely depleted and shown very little recovery, despite cod fishing moratoria being in place (Hutchings & Myers, 1994; Cardinale & Svedäng, 2004; FAO, 2020; ICES, 2021c–d). In the Skagerrak and Kattegat, current fisheries monitoring surveys catch almost no adult cod (Bland & Börjesson, 2020; Andersson *et al*., 2021). The juvenile abundances have also decreased in this area, but the trend has been less clear as the recruitment shows large interannual fluctuations (Svedäng, 2003; Cardinale & Svedäng, 2004). In addition, discard mortality of juvenile cod is still considered high, despite a landing obligation being in place since 2017 (ICES 2021c; Bryhn *et al*., 2022).

Genetic and tagging studies have identified two sympatric cod ecotypes residing in the North Sea-Skagerrak-Kattegat region (Knutsen *et al*., 2011; André *et al*., 2016; Barth *et al*., 2017; Svedäng *et al*., 2019). One of these ecotypes is genetically similar to North Sea cod and dominates in the offshore and outer coastal regions (Knutsen *et al*., 2018). This “offshore” ecotype consists, at least in part, of juvenile cod from the North Sea that are transported into the area with ocean currents (Stenseth, *et al.*, 2006; Jonsson *et al*., 2016). The influx of offshore juveniles varies between years, and is likely the main source of interannual variability in juvenile abundances (Knutsen *et al*., 2004; Stenseth *et al*., 2006). The offshore cod appear to utilise the Skagerrak and Kattegat coasts as a nursery for 2-4 years before migrating to the North Sea to spawn (Pihl & Ulmestrand, 1993; Svedäng *et al*., 2007; André *et al*., 2016; Hemmer-Hansen *et al*., 2020). The offshore ecotype is often referred to as “North Sea cod”, but, to date, it is unclear whether all fish make return migrations to the North Sea, or if some fraction completes its life cycle in offshore Skagerrak (Knutsen *et al*., 2018). The second ecotype, referred to as “fjord cod” in Norway and “coastal cod” in Sweden, is genetically similar to cod in the southern Kattegat and Öresund and is more common in the southern Kattegat and inshore coastal Skagerrak (Knutsen *et al*., 2018; Hemmer-Hansen *et al*., 2020). The coastal ecotype displays more resident behaviour and does not appear to undertake long-range migrations during the spawning season (Knutsen *et al*., 2011; Kristensen *et al*., 2021). The two ecotypes in the Skagerrak and Kattegat coexist on multiple spatial scales, but show differences in growth rate (Jørgensen *et al.* 2020), behaviour (Kristensen *et al.* 2021), and geographical distribution (Knutsen *et al.* 2018; Hemmer-Hansen *et al*., 2020). In addition, the ecotypes are genetically differentiated at both neutral (Knutsen *et al*., 2011) and potentially adaptive loci (Barth *et al*., 2019).

Cryptic population divergence evolving at small spatiotemporal scales is sometimes restricted to highly specific genomic regions. These genomic regions with above-average genetic differentiation may contain loci associated with adaptation to specific environmental conditions (Gagnaire *et al.*, 2015), or chromosomal rearrangements that restrict gene flow and maintain reproductive isolation between populations (Kirkpatrick & Barton, 2006). A striking trait of the Atlantic cod genome is the presence of four large chromosomal inversions on chromosome (chr) 1, 2, 7 and 12, which segregate in different populations across the species’ geographic range (Berg *et al*., 2016; Kess *et al*., 2020; Matschiner *et al.*, 2022). Loci within an inverted region of a chromosome may resist meiotic recombination in heterozygotes and are, in effect, often inherited as a single “supergene” (Dobzhansky & Epling, 1948). Hence, the alternative inversion states may represent combinations of alleles that are optimised for different environmental conditions (Wellenreuther & Bernatchez, 2018). As for the chromosomal inversions, differentiation in haemoglobin (Hb) genes is relatively well documented in cod. There are two main Hb components in cod, HbI and HbII (Sick, 1961), the former being the most well-studied. HbI genotype affects the oxygen affinity of Hb at different temperatures (Brix *et al*., 2004), which is reflected in the temperature preferences of cod with different HbI genotypes (Petersen & Steffensen, 2003). The putative roles of chromosomal inversions and Hb genes in the offshore-coastal ecotype divergence remain unknown.

Differentiated offshore and coastal ecotypes have also been documented for Atlantic cod in northern Norway (Michalsen *et al*., 2014), Iceland (Grabowski *et al*., 2011), Greenland (Pampoulie *et al*., 2011), and Canada (Ruzzante *et al*., 2000). In northern Norway, these ecotypes represent the migratory North-Eastern Arctic cod (NEAC; “skrei”) and the stationary Norwegian coastal cod (NCC; Michalsen *et al.*, 2014). The NEAC and NCC are strongly differentiated at all chromosomal inversions (Berg *et al*., 2016; Kess *et al*., 2019), as well as HbI genotype (Andersen *et al*., 2009). Recently, cod fisheries management in northern Norway has integrated genetic methods to assign individual cod to NEAC or NCC ecotype within 24 hours of capture, allowing a rapid regulation of the fisheries if catch proportions of the more sensitive NCC cod exceed a certain threshold (Dahle *et al*., 2018). In contrast, cod fisheries management in the Skagerrak and Kattegat does not account for the presence of two sympatric ecotypes in the area (ICES, 2020). Increasing the knowledge on the respective life history strategies and relative strengths of the offshore and coastal ecotypes has been highlighted as essential to the successful management of Skagerrak-Kattegat cod (Knutsen *et al*., 2018; Hemmer-Hansen *et al.*, 2020).

The low abundance of adult cod coupled with the variable juvenile recruitment in Swedish waters has led to hypotheses that most cod presently found along the Swedish Skagerrak coast have an offshore origin (Svedäng, 2003; Cardinale & Svedäng, 2004). However, recent studies in the Norwegian Skagerrak have shown high proportions of coastal ecotype cod inside Norwegian fjords (Jorde, Synnes *et al*., 2018; Knutsen *et al*., 2018), and potentially even fjord-specific ecotypes (Barth *et al*., 2019). Despite claims that most cod along the Swedish west coast have an offshore origin, the relative proportions of offshore and coastal ecotype cod have not been assessed explicitly. Biophysical models of larval drift suggest that spawning populations in the Kattegat and Öresund supply the majority of recruits to the Swedish west coast (Jonsson *et al*., 2016; Barth *et al*., 2017), while recent observations of early egg stages inside Swedish fjords suggest that local fjord spawning may still also occur (Svedäng *et al*., 2019). In addition to the lack of knowledge regarding the origin of recruits, recent declines in the offshore stocks in the North Sea (ICES, 2021a; 2021c) have raised questions about whether the Skagerrak-Kattegat coastal zone has lost some of its function as a nursery habitat for juvenile cod, regardless of ecotype. If true, this would provide an alternative explanation for the declines in adult cod abundance along the Skagerrak-Kattegat coast.

In this study, we genotyped juvenile cod collected in the Skagerrak, Kattegat and Öresund in 2019 and 2020, using both a panel of targeted SNP loci developed to assign individuals to ecotype and sex, and a larger panel of genome-wide SNP loci obtained through 2b-RAD sequencing. The aims were: 1) to examine whether there are juvenile cod of both offshore and coastal origins along the Swedish west coast, and, if so, if there is geographical and interannual variation in the juvenile recruitment of both ecotypes; 2) to analyse whether the ecotypes are genetically differentiated at candidate loci associated with environmental adaptation; and 3) to explore potential genetic substructure within the coastal ecotype.

## Methods

### Sampling

Juvenile cod were collected along the Swedish west coast in 2019 and 2020, and along the Norwegian Skagerrak coast in 2020. Adult cod from the North Sea, Skagerrak, Kattegat, and Öresund were also included as reference individuals (Figure 1). The fishing gear used included bottom trawls, beach seines, fyke nets, shrimp trawls, lobster traps, and cages (Table S1). Fin clips were taken and stored in 95% ethanol at −20° C. DNA was extracted from the tissue samples with a Qiagen DNeasy^®^ Blood & Tissue kit following the manufacturer’s protocol. To allow inter-and intra-cohort analyses, juvenile samples that had not already been aged from otolith readings were assigned to cohort based on their total length (TL). Informed by length distribution graphs of both ecotypes (Figure S1), we assigned individuals with a TL ≤ 150 mm as 0-group cod and those with a TL > 150 mm as 1-group.

**Figure 1.**
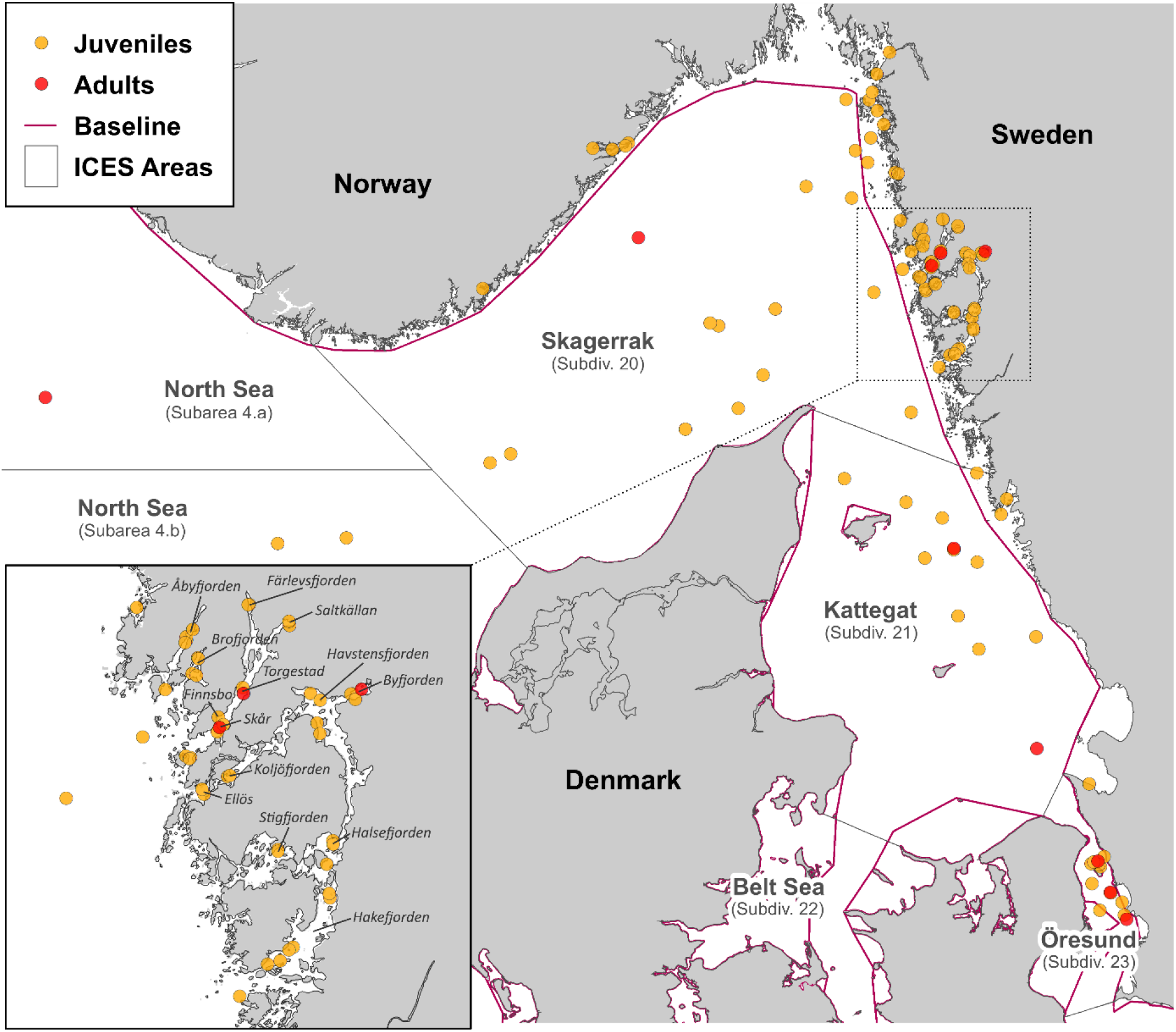
Map of the study area. Sampling stations are indicated with points, with colour indicating the type of sample (juvenile/adult). The national baselines of Norway, Sweden, and Denmark, which define the outer coastline, are indicated with purple lines. ICES subareas and subdivisions are indicated with narrow black lines.

### Targeted loci

A total of 1002 juvenile and adult individuals were genotyped using a set of 65 single nucleotide polymorphism (SNP) loci (Table S2). Of these, 40 loci were selected for discriminating between the offshore and coastal ecotypes along the Norwegian Skagerrak coast. These 40 loci were chosen from a panel of > 9000 SNP loci used to genotype cod in the Norwegian Skagerrak (see Jorde, Kleiven *et al*., 2018). Loci were ranked by the level of genetic divergence (Nei’s *G*_ST_) between the offshore and coastal ecotypes, excluding closely linked loci (composite linkage disequilibrium, CLD > 0.5). Of these 40 loci, 17 have been included in previous studies to discriminate between the ecotypes (Jorde, Kleiven *et al*., 2018; Jorde, Synnes *et al*., 2018; Knutsen *et al*., 2018). In addition, we included 8 sex-diagnostic loci (Star *et al*., 2016), and 19 candidate loci, putatively involved in environmental adaptation: 13 loci located within chromosomal inversions - 3 loci each on chr 1, 2, and 7, and 4 loci on chr 12 (Berg *et al*., 2015; Kess *et al*., 2020) - and one SNP locus within each Hb subunit in the *β5-α1-β1-α4* globin gene cluster on chr 2 (Borza *et al*., 2010). SNP genotyping was performed on an Agena MassARRAY Platform, as described by Gabriel *et al.* (2009), at the Mutation Analysis core Facility at Karolinska Institutet, Huddinge, Sweden. After quality control and filtering (see supplementary note S1), 52 SNP loci remained; 33 from the ecotype-diagnostic panel, 5 sex-linked loci, 10 loci located within inversions (2 each on chr 1 and 2, and 3 each on chr 7 and 12), and 4 Hb loci. The final dataset consisted of 406 and 488 juveniles collected in Swedish waters in 2019 and 2020, respectively, 25 juveniles from the Norwegian fjords, along with 11 adult reference cod from the North Sea, 6 from Norwegian Skagerrak, 18 from Gullmarsfjorden, 3 from Byfjorden, 17 from Kattegat, and 13 from Öresund (Figure 1).

#### Ecotype assignment

We assigned all individuals to ecotypes through K-means clustering using adegenet (v2.1.3; Jombart, 2008). Given that our panel was developed to distinguish the offshore and coastal ecotypes, we performed the K-means clustering assuming two groups (K = 2), corresponding to the two ecotypes. Individuals were assigned to the offshore or coastal ecotype according to whether they were assigned to the same group as adult reference individuals from the North Sea or from the Kattegat and Öresund, respectively. The ecotype assignment thus resulted in 491 offshore and 496 coastal ecotype individuals. The assignment was then evaluated using four different methods (see supplementary note S2): discriminant analysis of principal components (DAPC; adegenet), principal component analysis (PCA; ade4 v1.7-16; Dray & Defour, 2007), sparse non-negative matrix factorization (SNMF; LEA v3.1.2; Frichot & François, 2015), and population assignment (assignPOP v1.2.2; Chen *et al*., 2020). Lastly, we estimated the genetic divergence between the ecotypes (*F*_ST_) and its significance, using a permutation test (wcFst; 10 000 permutation replicates) in strataG (v2.5.01; Archer *et al*., 2016). We used mapplots (v1.5.1; Gerritsen, 2018) to visualise the geographical distribution of the ecotypes.

#### Assignment of sex and inversion state

For both the sex- and inversion-linked SNP loci, PCA revealed clear clustering of individual genotypes along PC1 (Figure S2). Hence, we used the individual PC1 coordinates to assign sex and inversion state to each individual, similar to the methods described by Ma & Amos (2012). To determine which inversion state was the ancestral and which was the derived for each chromosome, we assumed that the most frequent rearrangements in our study corresponded to those reported for Norwegian coastal cod and Western Baltic cod by Matschiner *et al.* (2022). Note that different studies have used different nomenclature for the alternative inversion states, see Table S3. Individuals with one homozygous and one heterozygous SNP locus within the same inversion were removed from downstream analyses (chr 1: n = 31, chr 2: n = 5, chr 12: n = 16). The four Hb loci were analysed individually.

#### Interannual differences in ecotype proportions

We tested for inter- and intra-cohort differences in ecotype proportions between years using weighted analyses of variance (ANOVA), as applied in stats (v4.1.2; R Core Team, 2021). For the inter-cohort analysis, we only included stations at which 0-group cod were collected in both years, whilst for the intra-cohort analysis, we only included stations at which juveniles from the 2019 cohort were collected in both 2019 and 2020 (as 0- and 1-group cod, respectively). In both cases, station-wise estimates of ecotype proportions were weighted against the number of individuals included from each station and year. Weighted ANOVAs were then performed separately for offshore and inshore stations. Öresund stations were excluded from this analysis, as the sample sizes in 2020 were very low.

#### Genetic differences between ecotypes at candidate loci

We tested for differences in allele frequencies between ecotypes for the inversion- and Hb loci by calculating locus-wise *F*_ST_ between the ecotypes using strataG (10 000 permutation replicates). The assigned inversion state of each individual was recoded as a single superlocus genotype for this analysis, to avoid biasing the *F*_ST_ estimates by including multiple non-independent markers. To test for differences in sex ratio, we performed Pearson’s χ^2^ test with Yates’ continuity correction, as applied in STATS.

#### Environmental predictors of ecotype and candidate locus genotype

We used regression modelling and model selection to explore potential geographical correlates with station-wise ecotype proportions and sex ratios, and with individual genotypes at candidate loci. We only included 0-group juveniles from 2019 and 2020 collected in Skagerrak (ICES Subdivision 20), Kattegat (ICES Subdivision 21), and Öresund (ICES Subdivision 23) in these analyses. This was done to avoid including age classes with small sample sizes per station and subdivisions sampled at very few stations. Hence, these analyses excluded all 1-group juveniles and adults, as well as all individuals collected in the North Sea.

For the ecotype proportions and sex ratios we fitted weighted linear regression models using stats. We included year and ICES Subdivision as categorical explanatory variables, bottom depth and distance inshore from the coastal baseline for each station as linear explanatory variables, and the number of individuals per station as weights. ICES Subdivision was included as an explanatory variable, as the subdivisions correlate with the current ICES advisory units for cod in the area: the “North Sea, eastern English Channel, Skagerrak” advisory unit includes Subdivision 20; the “Kattegat” advisory unit corresponds to Subdivision 21; and the “western Baltic Sea” advisory unit includes Subdivision 23 (ICES, 2020). Distance from the baseline (positive value inshore, negative value offshore) and depth were included as proxies for spatial environmental variation, e.g., in temperature, salinity, or oxygen conditions. For the individual chromosomal inversion state, and genotypes at Hb loci, we fitted ordinal logistic regression models using MASS (v7.3-54; Venables & Ripley, 2002). As the ordinal regression models were fitted against individual-level data, we included the ecotype and sex of each individual as categorical explanatory variables, in addition to year, ICES Subdivision, depth and distance from the baseline. Depth was log-transformed to address the skewed distribution of this variable, and to approach a linear relationship with the response variables. For more details, see supplementary note S3.

We applied a model selection approach to the full suite of models to identify the most informative predictors. The quality of each model was determined by calculating the sample-size-corrected Akaike’s information criterion (AICc), using AICcmodavg (v2.3-1; Mazerolle, 2020). Models with an AICc difference (ΔAICc) < 2 were treated as having equal levels of support, following Burnham & Anderson (2004), and are referred to as the “top-ranked” models henceforth. We consider our analyses of environmental correlates as exploratory (see Tredennick *et al*., 2021), and thus, we opted against selecting a single model to make statistical inferences from. Instead, we have identified the most important explanatory variables across the top-ranked models and their respective effects, to indicate the main geographical correlations (see Burnham & Anderson, 2004; Tredennick *et al*., 2021).

### Genome-wide loci

A subset of 268 individuals was selected for genome-wide genotyping using 2b-RAD sequencing. The subset included juveniles from Swedish (n=181) and Norwegian (n=12) Skagerrak fjords, the Kattegat (n=18), and Öresund (n=16), together with adult reference cod from the North Sea (n=10), Gullmarsfjorden (n=9), Byfjorden (n=3), Kattegat (n=10) and Öresund (n=9). The main aim of this approach was to analyse potential substructure within the coastal ecotype, and specifically if juveniles collected in fjords were distinct from cod in the Kattegat. However, we also included juveniles assigned to the offshore ecotype, which enabled comparisons of any potential coastal substructure to the already described offshore-coastal ecotype divergence. It also allowed for independent evaluation of the ecotype assignment using the ecotype-diagnostic SNP panel (see supplementary note S4).

#### Library preparation and sequencing

The DNA integrity was assessed in a 1% agarose gel and the quality using a NanoDrop^®^ ND-1000 spectrophotometer. The quality of the DNA extractions was improved with a purification step using a Zymo DNA Clean & Concentrator™-25 Kit. The 2b-RAD libraries (Wang *et al*., 2012) were prepared following a modified protocol by Mikhail Matz (https://github.com/z0on/2bRAD_GATK/blob/master/2bRAD_protocol_june1_2018.pdf) as described in Kinnby *et al.* (2020).

#### Mapping and filtering

Trimming, filtering, and genotype calling was done following Pereyra *et al.* (2022). Trimmed and quality-filtered sequences were mapped to the cod genome assembly gadMor3.0 (NCBI Bioproject accession no. <u>PRJEB33455</u>).

#### Genotype calling

Individual RAD libraries produced 0.3–14.8 (median: 4.4) million reads per individual. Mapping to the gadMor3.0 genome assembly resulted in alignment rates ranging 74.9-95.9 % (median 94.0 %). Technical replicates rendered 15 320 SNPs that were used as “true” SNP dataset for recalibration of variants. A total of 55 449 SNPs were called, and a set of 23 364 SNP loci and 235 individuals was obtained after filtering following Pereyra *et al.* (2022). Subsequent removal of loci with a minor allele frequency < 5 % resulted in 9956 SNP loci for downstream analyses. A complete list of the filtered loci is provided in Table S4.

#### Inversion scans

To identify loci located within inverted chromosomal regions, the filtered SNPs were scanned for chromosomal inversions using inveRsion (v1.40.0, Cáceres, 2021). For details, see supplementary note S5.

#### Local coastal populations

We performed PCAs in adegenet to visualise the genetic distances between coastal ecotype individuals at genome-wide loci located outside of inversions. We also calculated pairwise multi-locus *F*_ST_ between sampling sites with strataG (1000 permutation replicates). For stations with small sample sizes, individuals from multiple nearby stations were pooled, thus representing the juvenile assemblage of a larger area (Table S5). Due to low 2b-RAD genotyping success for the fjord reference samples, adults from Gullmarsfjorden and Byfjorden were also pooled. The dimensions of the pairwise *F*_ST_ matrix were then reduced using multi-dimensional scaling (MDS) in STATS.

#### Outlier analyses

Pairwise outlier tests were performed using BayeScan (v2.1; Foll & Gaggiotti, 2008) and OutFlank (v0.2; Whitlock & Lotterhos, 2014) with default settings, to identify loci displaying patterns of non-neutral selection between the ecotypes. Annotation of outlier loci was performed by running 2.5 kb flanking regions of each outlier locus through blastx, to match to all non-redundant GenBank CDS translations+PDB+SwissProt+PIR+PRF databases. Loci with annotated hits were subsequently searched for Gene Ontology (GO) terms using PANTHER DB (Mi *et al.*, 2021) and PANTHER’s tool (Thomas *et al.*, 2006) to access the list of GO annotations (Ashburner *et al.*, 2000; Gene Ontology Consortium, 2021).

Unless else stated, all data filtering and statistical analyses were performed in R (v4.1.2; R Core Team, 2021), using RStudio (v1.4.1717; RStudio Team, 2021). All p-values were corrected for multiple testing with stats, using the false discovery rate (FDR) method with the threshold for significance set at q < 0.05.

## Results

### Ecotype assignment

The DAPC (Figure 2A) and PCA (Figure 2B) performed with the ecotype-diagnostic SNP panel separated the ecotypes into two distinct clusters. The two ecotypes were also separated to a large extent in the PCA using genome-wide loci located outside of inverted chromosomal regions (Figure 2C). The latter PCA was largely unchanged when outlier loci (Figure 3) were removed (Figure 2D-E). The ecotype assignment was further corroborated by alternative assignment methods (Supplementary note S2; Figure S3; Table S6). Multi-locus *F*_ST_ between the ecotypes was 0.14 (p < 0.01) for the ecotype-diagnostic loci, and 0.006 (p < 0.01) for the 2b-RAD genome-wide loci outside of inverted chromosomal regions.

**Figure 2.**
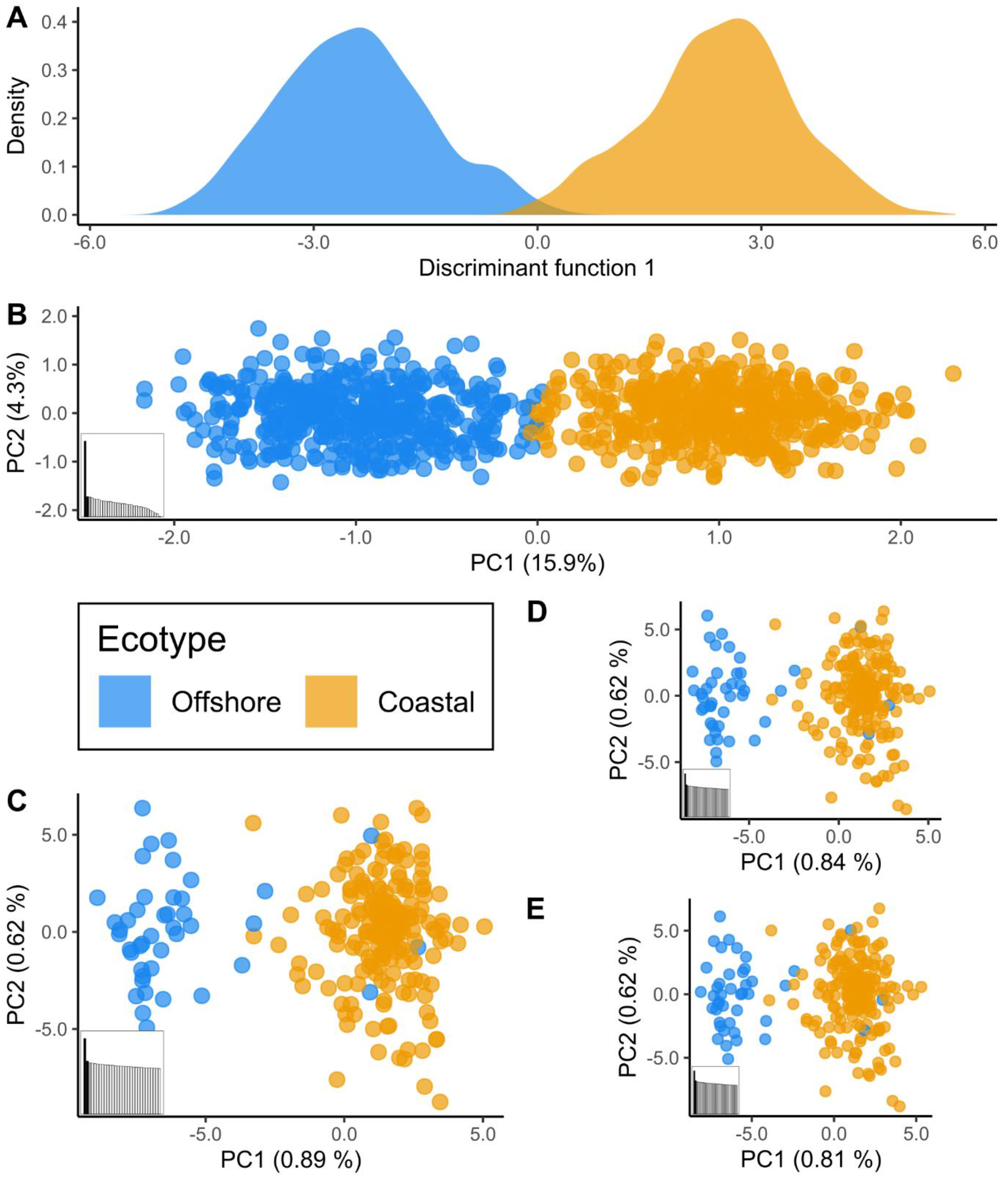
**A)** DAPC scores along discriminant function 1 and **B)** PCA score plot for all 987 individuals, based on genotypes at the 33 ecotype-diagnostic loci. Subplots **C-E)** show PCA score plots based on the genome-wide loci located outside of inversions, with **C)** showing all these loci, **D)** excluding BayeScan outliers, and **E)** excluding OutFLANK outliers. Colour corresponds to the assigned ecotype of each individual, according to the ecotype-diagnostic SNP panel. The aspect ratio between PC1 and 2 is scaled against the relative proportions of variance explained by each PC (in parentheses).

### Geographical distribution of ecotypes

Juveniles (0- and 1-group) of the two ecotypes co-occurred both offshore and inshore across the sampled region (Figure 4), but the ecotypes also showed differences in their geographical distribution that were highly stable between the years. Juveniles in offshore areas and the northern Skagerrak predominantly assigned to the offshore ecotype, whilst juveniles in inshore areas, southern Kattegat, and Öresund predominantly assigned to the coastal ecotype. With few exceptions, the proportion of coastal ecotype juveniles generally increased toward the innermost parts of the fjords, although the offshore ecotype was also found in this environment (Figure 4, inset maps).

**Figure 3.**
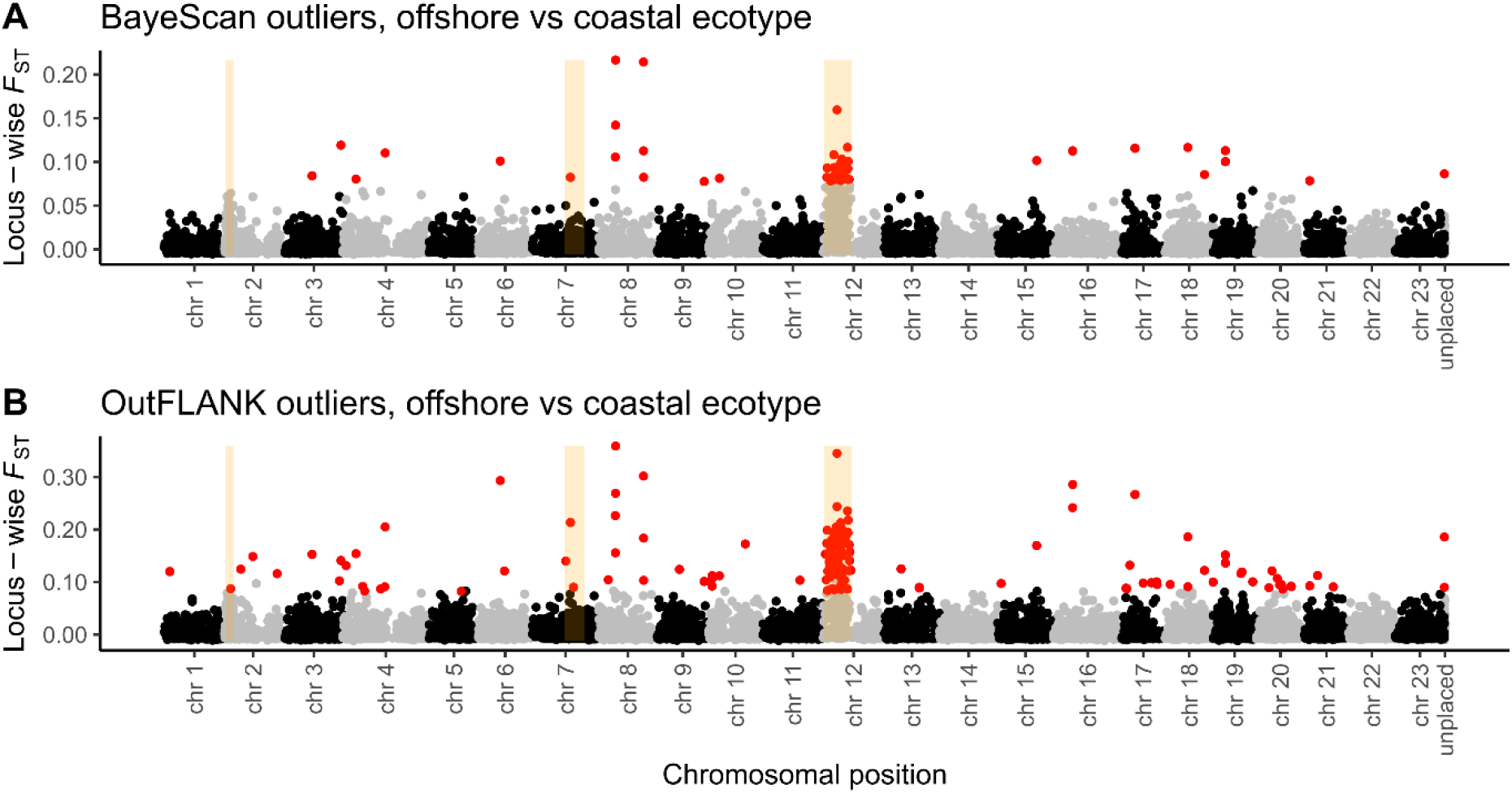
Manhattan plots of pairwise *F*_ST_ between the two ecotypes for all 2b-RAD SNP loci, with outlier loci detected by **A)** BayeScan and **B)** OutFLANK indicated as red points. Note that the *F*_ST_ values shown in **B)** are not corrected for sample size differences, as these are the input values for the OutFLANK algorithm. The inverted regions detected by the inveRsion scans are highlighted in orange.

**Figure 4.**
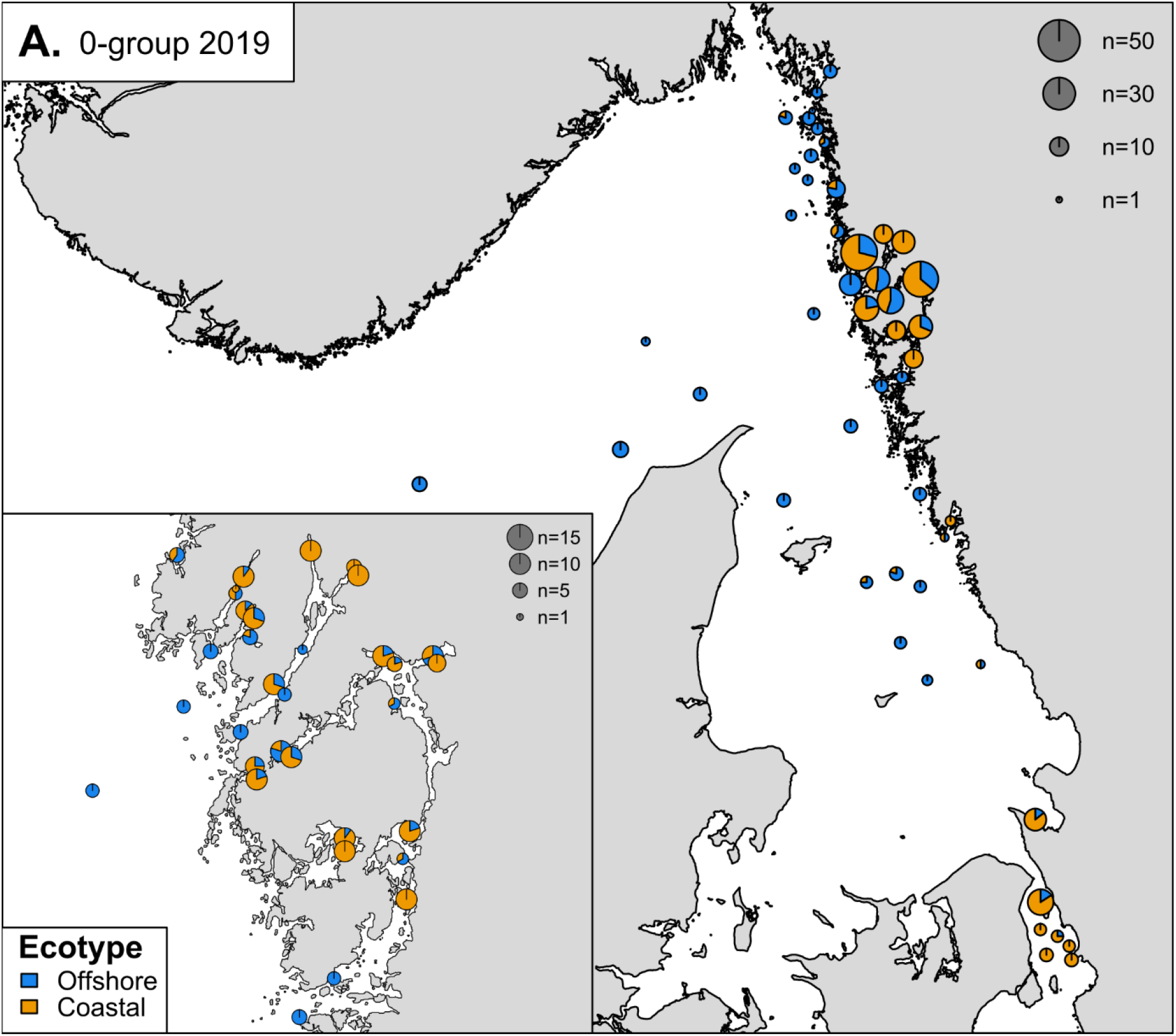
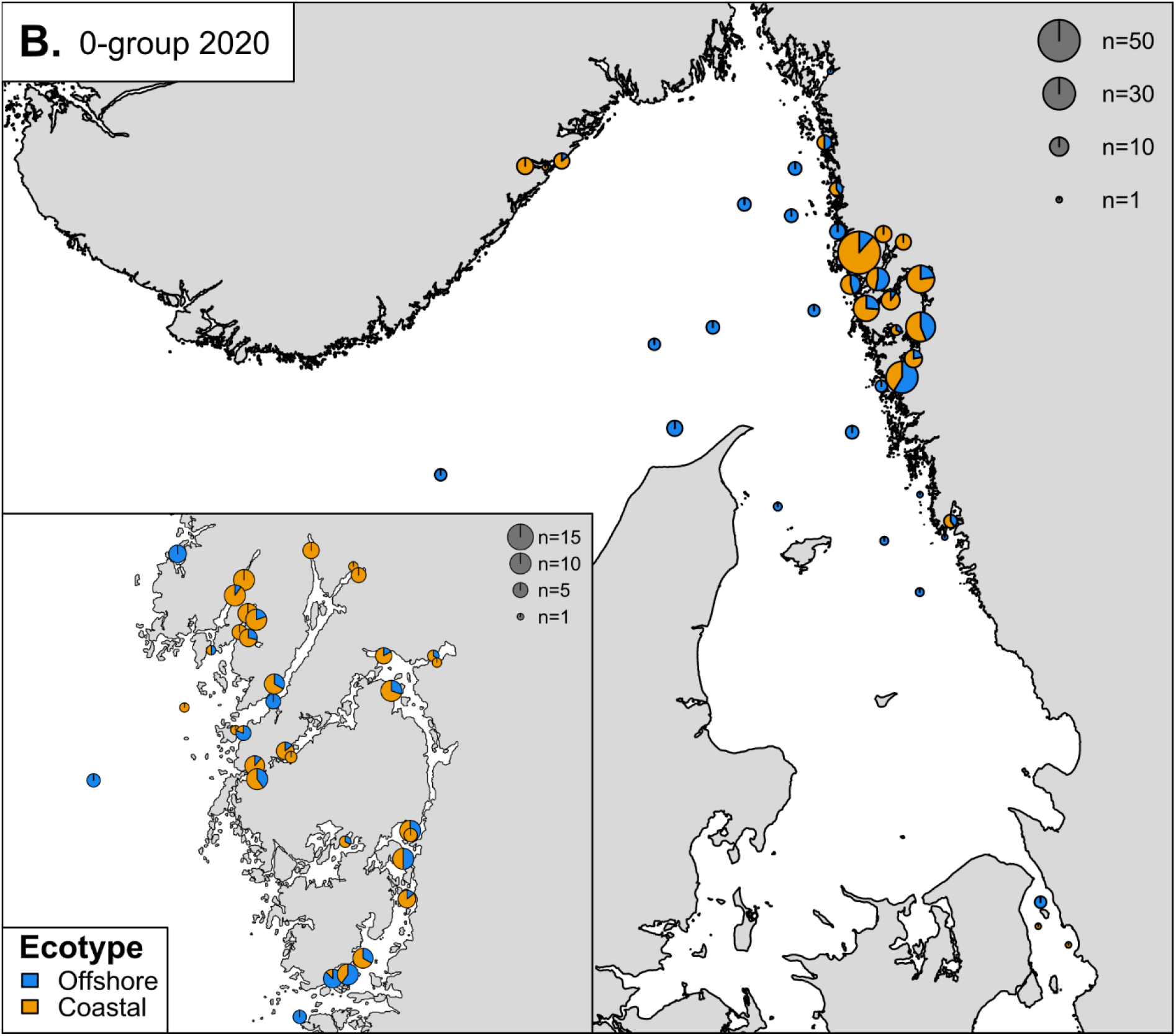
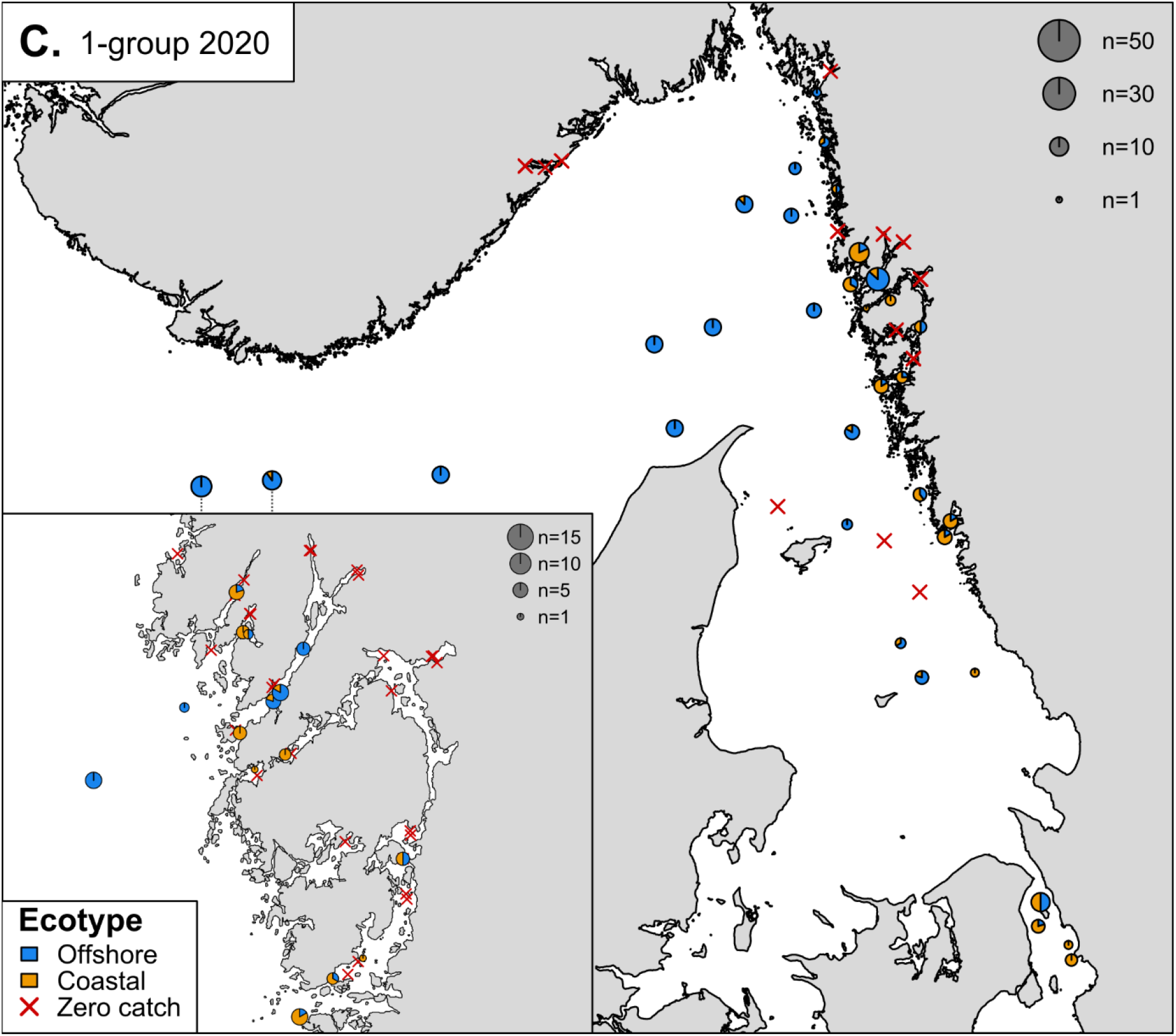
Maps of ecotype distributions for **A)** 0-group cod in 2019, **B)** 0-group cod in 2020, and **C)** 1-group cod in 2020. The area of each pie indicates the number of individuals per station, and the size scale is indicated with grey pies. Note that different size scales are used for full maps and inset maps, but that the same scales are used in **A, B**, and **C**. The size of each pie slice indicates the proportion of the offshore (blue) or coastal ecotype (orange). Sampled stations at which no 1 -group cod were caught in 2020 are indicated with red crosses in **C**.

The ecotype proportions for 0-group cod were very similar in 2019 and 2020 (Figure 4A vs 4B), at both offshore (F_1,17_ = 3.04, q = 0.20) and coastal stations (F_1,67_ = 0.48, q = 0.49). Ecotype proportions for 0-group in 2019 and 1-group cod in 2020 (intra-cohort) were also similar at offshore stations (F_1,12_ = 0.91, q = 0.48), but differed at coastal stations (F_1,30_ = 7.33, q = 0.04). This was due to an increase in the proportion of coastal ecotype in the outer archipelago between 2019 (0-group, Figure 4A) and 2020 (1-group, Figure 4C). However, 1-group cod were absent at many stations in 2020, especially in inshore Skagerrak.

### Differentiation at candidate loci

The two ecotypes showed differentiation in several genomic regions that have previously been linked with environmental adaptation (Figure 5). These included inversions on chr 2 (*F*_ST_ = 0.08, q < 0.01), chr 7 (*F*_ST_ = 0.03, q < 0.01), and chr 12 (*F*_ST_ = 0.21, q < 0.01), as well as the Hb-β1 (*F*_ST_ = 0.01, q < 0.01) and Hb-β5 loci (*F*_ST_ = 0.02, q < 0.01). However, the ancestral chr 1 inversion state appeared to be fixed in both ecotypes, and the ecotypes did not differ in genotype frequencies at Hb-α1 (*F*_ST_ = 0.00, q = 1.00) and Hb-α4 (*F*_ST_ = 0.00, q = 0.11), or sex ratio (χ^2^ = 0.37, df = 1, q = 0.62).

**Figure 5.**
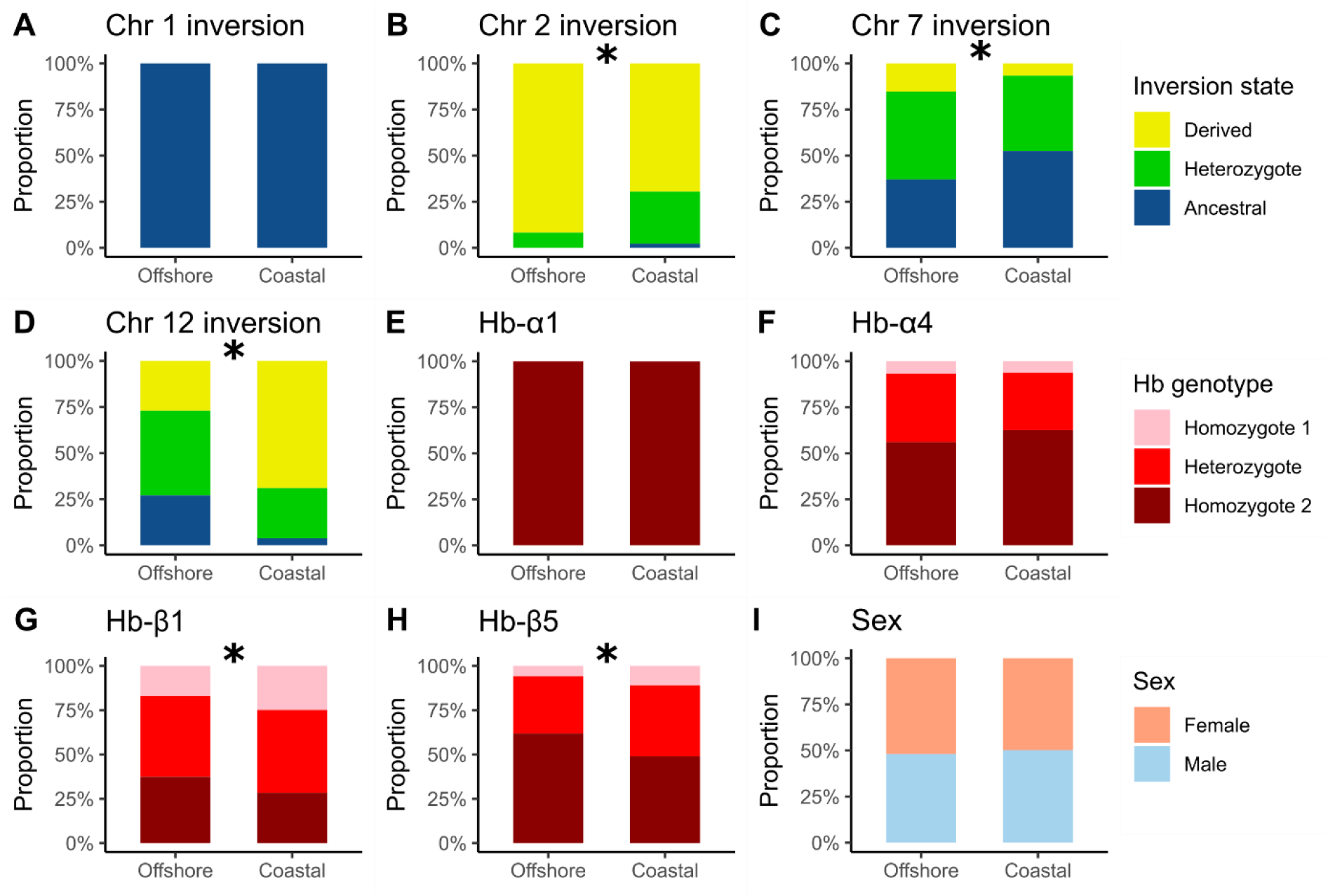
Barplots showing the overall **A-D)** inversion state frequencies, **E-H)** genotype frequencies at haemoglobin (Hb) loci, and **I)** sex ratio for each ecotype. Asterisks indicate significant *F*_ST_ between the ecotypes after FDR-correction for multiple testing.

### Environmental predictors

#### Ecotype

Our exploration of potential drivers behind the distribution of ecotypes using model selection returned 23 models in total (Table 1), with three models of equal quality (ΔAIC < 2). The simplest of the three models, Model 1, included only ICES Subdivision and depth in the predictor, and both had significant effects (Table S7). Subdivision and depth were included in all the top-7 models, and depth was included in all top-13 models. Models 2 and 3 also included these variables, and the additional explanatory variables included in these models all had non-significant effects. According to all three top-ranked (ΔAICc < 2) models, the proportion of coastal ecotype was predicted to be similar between the Skagerrak and Kattegat, but higher in the Öresund, and to decrease with depth in all subdivisions.

**Table 1.**
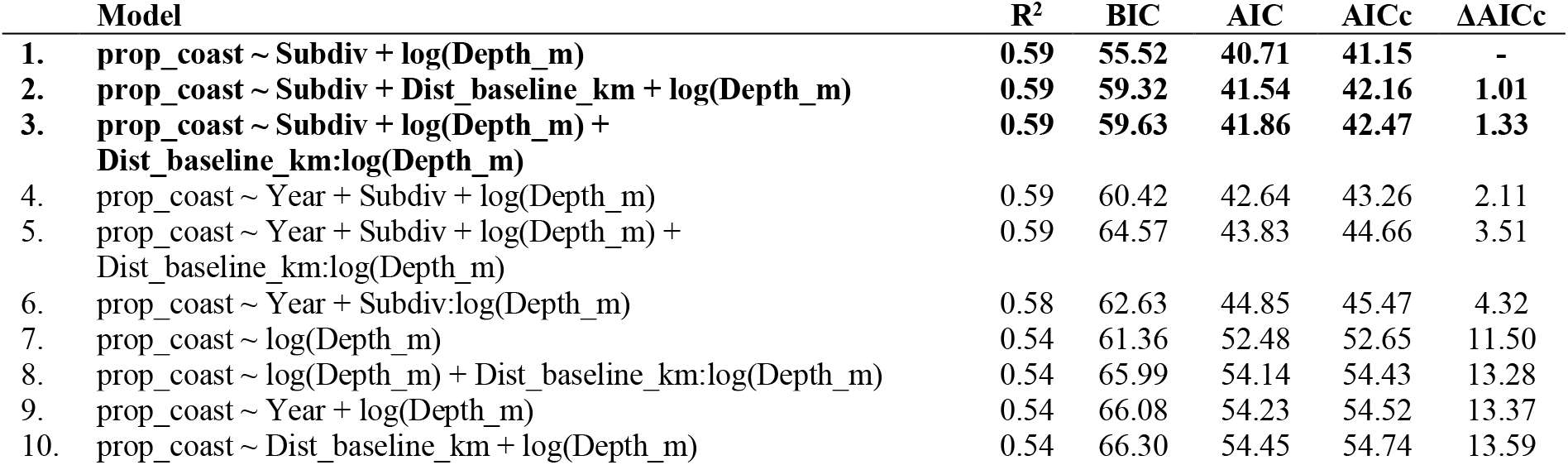
AIC table for the 10 highest-ranked linear regression models fitted against the station-wise proportion of coastal 0-group juveniles. The top-ranked models (ΔAICc < 2) are indicated in bold.

#### Candidate locus genotype

Ecotype was the single most important explanatory variable for most loci. However, the chromosomal inversions on chr 2, 7 and 12 were also correlated with environmental variables. For the chr 2 inversion, both distance inshore from the baseline and ecotype were important explanatory variables (Table 2). Both variables were included in all 4 top-ranked models and had significant effects, whilst all additional explanatory variables in these models were non-significant (Table S9). In all top-ranked models, the ancestral chr 2 inversion state was more common in the coastal ecotype, and increased in frequency with distance from the baseline. Model selection for the chr 7 inversion returned 6 top-ranked models, for which the only common explanatory variable was bottom depth, either alone or as an interaction effect with ecotype (Table 3). A Subdivision × distance interaction was also present in some of these models. Thus, it appears that depth was the most important explanatory variable for the chr 7 inversion, but its effect may be moderated by ecotype, Subdivision, and distance from the baseline. Overall, the frequency of the derived inversion state increased with depth, but this effect was stronger in the offshore ecotype. Some models suggested that the derived state also decreased in frequency with distance from the baseline, however only significantly in the Kattegat or the offshore ecotype (Table S10). For the chr 12 inversion, all 8 top-ranked models included various combinations of ecotype, depth, and distance from the baseline as predictors (Table 4). The top-ranked model with the fewest explanatory variables (model #5) included distance and the depth × ecotype interaction, the latter having the strongest effect. A distance × depth interaction was also frequent across the top-ranked models. Shared among the top-ranked models was that the ancestral allele increased sharply with depth in the offshore ecotype, but not in the coastal ecotype, whilst distance had a very small effect (Table S11).

**Table 2.**
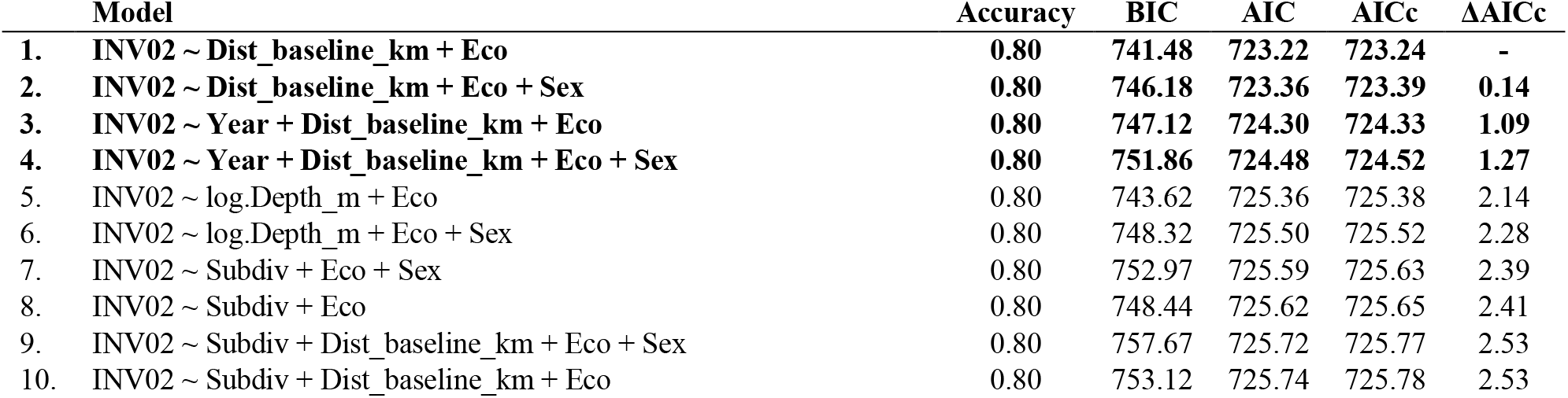
AIC table for the 10 highest-ranked ordinal logistic regression models fitted against the individual chr 2 inversion state genotypes for 0-group juveniles. The top-ranked models (ΔAICc < 2) are indicated in bold. “Accuracy” refers to the proportion of correctly assigned genotypes for each model.

**Table 3.** AIC table for the 10 highest-ranked ordinal logistic regression models fitted against the individual chr 7 inversion state genotypes for all 0-group juveniles. The top-ranked models (ΔAICc < 2) are indicated in bold. “Accuracy” refers to the proportion of correctly assigned genotypes for each model.

| | Model | Accuracy | BIC | AIC | AICc | $\Delta\text{AICc}$ |
| --- | --- | --- | --- | --- | --- | --- |
| 1. | **INV07 ~ Subdiv:Dist_baseline_km + log.Depth_m:Eco** | **0.54** | **1342.54** | **1310.57** | **1310.62** | - |
| 2. | **INV07 ~ log.Depth_m + Subdiv:Dist_baseline_km** | **0.53** | **1338.56** | **1311.15** | **1311.19** | **0.57** |
| 3. | **INV07 ~ log.Depth_m + Dist_baseline_km:Eco** | **0.52** | **1334.91** | **1312.07** | **1312.10** | **1.48** |
| 4. | **INV07 ~ log.Depth_m:Eco** | **0.53** | **1330.55** | **1312.28** | **1312.30** | **1.68** |
| 5. | **INV07 ~ Year + Dist_baseline_km + Subdiv:Dist_baseline_km + log.Depth_m:Eco** | **0.53** | **1348.94** | **1312.39** | **1312.46** | **1.84** |
| 6. | **INV07 ~ Sex + Subdiv:Dist_baseline_km + log.Depth_m:Eco** | **0.55** | **1349.00** | **1312.46** | **1312.53** | **1.90** |
| 7. | INV07 ~ Dist_baseline_km + log.Depth_m:Eco | 0.53 | 1335.83 | 1312.99 | 1313.02 | 2.40 |
| 8. | INV07 ~ Year + log.Depth_m + Subdiv:Dist_baseline_km | 0.53 | 1344.95 | 1312.97 | 1313.03 | 2.40 |
| 9. | INV07 ~ log.Depth_m + Sex + Subdiv:Dist_baseline_km | 0.53 | 1345.03 | 1313.06 | 1313.11 | 2.49 |
| 10. | INV07 ~ Subdiv:log.Depth_m + log.Depth_m:Eco | 0.54 | 1340.66 | 1313.25 | 1313.29 | 2.67 |

**Table 4.**
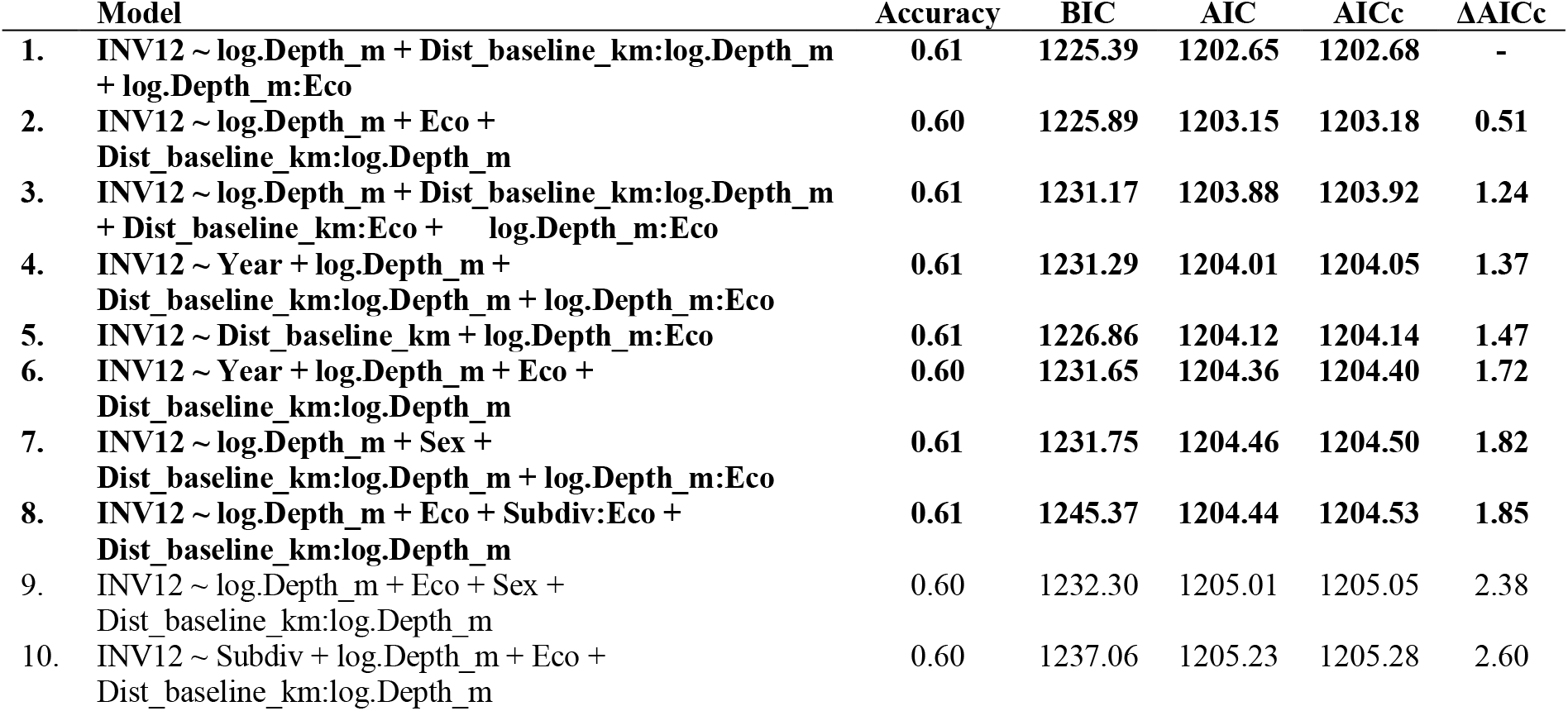
AIC table for the 10 highest-ranked ordinal logistic regression models fitted against the individual chr 12 inversion state genotypes for all 0-group juveniles. The top-ranked models (ΔAICc < 2) *are* indicated in bold. “Accuracy” refers to the proportion of correctly assigned genotypes for each model.

We excluded the chr 1 inversion and Hb-α1 loci from the regression analyses as the total genetic variance was too low. For the Hb-α4, Hb-β1 and Hb-β5 loci, the models including only ecotype as predictor were among the top-ranked models, and no other variables appeared to be important in predicting the genotypes at these loci (Table S12-S14). In addition, the intercept-only model was among the top-ranked models for sex ratio (Table S8), indicating that sex ratio of the 0-group juveniles did not correlate with any of the included explanatory variables.

### Outlier analyses

From the genome-wide SNP panel, BayeScan and OutFLANK identified 57 and 156 outlier loci between the ecotypes, respectively (Figure 3). All 57 outlier loci detected by BayeScan were also detected by OutFLANK, and more than half of the outlier loci were located within the inverted region on chr 12, for both methods (33/57 for BayeScan and 80/156 for OutFLANK). Gene annotation of all 156 outlier loci resulted in 80 unique gene hits obtained for 85 loci (see Table S15), and GO terms were available for 71 of these genes (see Table S16). However, no enrichment analysis was performed on these GO terms, as the number of genes was too small.

### Local coastal populations

When exploring putative genetic substructure within the coastal ecotype, the PCA based on 9418 genome-wide loci outside inversions revealed no distinct clusters (Figure 6A). Instead, there was a high degree of overlap among stations, and each PC explained similarly low proportions of the total genetic variation within the coastal ecotype (< 1 %). Pairwise *F*_ST_ values (Table 5) were significant between the North Sea offshore adults and most of the coastal samples (*F*_ST_ = 0.006 – 0.010, q = 0.008 – 0.014), but there were also significant *F*_ST_ values within the coastal ecotype, between juveniles from Norwegian fjords (Risør area) and seven of the innermost Swedish fjords – Brofjorden, Färlevsfjorden, Saltkällan, Ellösefjorden, Havstensfjorden, Byfjorden, and Hakefjorden (*F*_ST_ = 0.003 – 0.004, q = 0.008 – 0.045). However, *F*_ST_ was non-significant for all coastal ecotype samples compared to both Kattegat and Öresund spawning adults, and only significant between two Swedish fjord samples – the two adjacent fjords Havstensfjorden and Byfjorden (*F*_ST_ = 0.003, q = 0.040). The MDS plot based on these pairwise *F*_ST_ values (Figure 6B), similarly to the PCA, showed a large cluster clearly separated from the North Sea adults. Within this cluster, coastal ecotype juveniles from the Skagerrak, Kattegat and Öresund clustered together with spawning adults from both Kattegat and Öresund. In contrast to the PCA score plot, coastal ecotype juveniles from Norwegian fjords were separated from the main coastal cluster, indicating genetic divergence from the Swedish samples.

**Figure 6.**
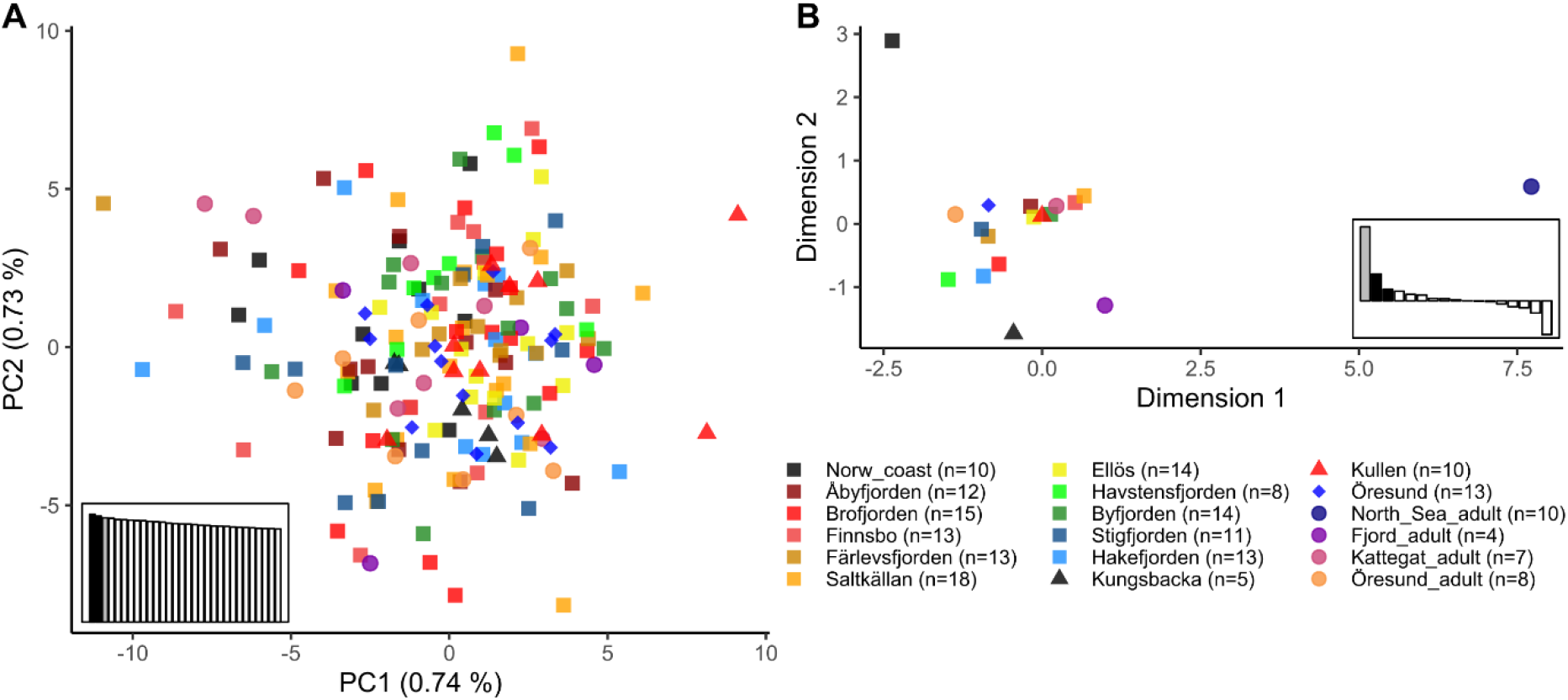
**A)** PCA score plot showing the genotypic distances between individuals assigned to the coastal ecotype, and **B)** MDS plot based on the *F*_ST_ values in Table 4. Note that **A** only includes individuals assigned to the coastal ecotype, whereas **B** also includes the North Sea adults as an outgroup. Genetic distances in both plots are based on the genome-wide loci located outside of inverted regions. The scree plots show the relative eigenvalues of **A)** the first 30 PCs, or **B)** all MDS dimensions. Squares = Skagerrak juveniles, triangles = Kattegat juveniles, diamonds = Öresund juveniles, and circles = adults.

**Table 5.**
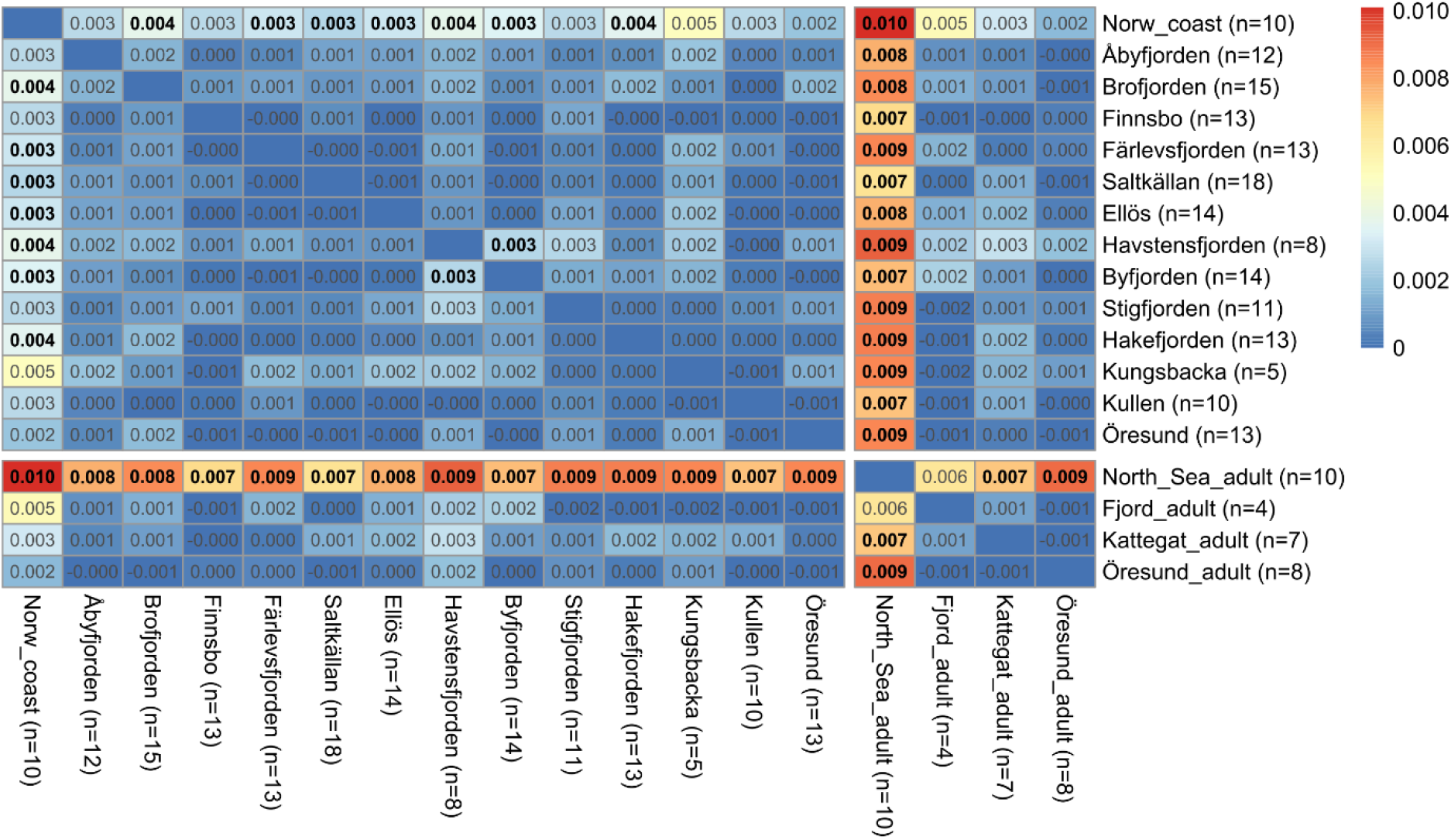
Pairwise *F*_ST_ between stations, including only coastal ecotype individuals from each station, and North Sea adults as an outgroup. The *F*_ST_ values are based on the genome-wide loci located outside of inversions, and significant values after FDR-correction (q < 0.05) are indicated in bold font. Individuals have been grouped based on their sampling station, irrespective of survey and year. The total number of individuals included per station is given within parentheses. The four adult populations are located at the bottom and right-hand side of the table.

## Discussion

Our results show that the 2019 and 2020 cohorts of Atlantic cod collected along the Swedish west coast were mechanical mixtures of offshore and coastal ecotype juveniles. Furthermore, the coastal ecotype was dominant in many locations where both ecotypes co-occurred, especially in the inshore and southern regions. This contrasts with the hypothesis that juvenile cod found along the Swedish Skagerrak coast mainly originate from offshore spawning areas in the North Sea or Skagerrak (Svedäng, 2003; Cardinale & Svedäng, 2004). In addition to differences in geographical distribution between the ecotypes, the ecotypes were differentiated at multiple SNP loci that may be involved in adaptation to local environmental conditions.

### Ecotype distribution

The geographical distribution of the two ecotypes in Swedish waters was consistent with previous studies, showing a dominance of the offshore ecotype in offshore Skagerrak, whereas the coastal ecotype dominates inshore localities in the Skagerrak, southern Kattegat, and Öresund (Barth *et al.*, 2017; Knutsen *et al*., 2018; Hemmer-Hansen *et al*., 2020). However, the present study provides the first fine-scaled overview of the geographical distribution of both ecotypes in Swedish waters. Juveniles from both ecotypes coexist in coastal areas at very small spatial scales, as has been described also in Norwegian fjords (Knutsen *et al*., 2018; Jorde, Synnes *et al*., 2018). The geographical distribution of ecotypes was highly stable between 2019 and 2020, which indicates a low interannual variability in the spatial recruitment patterns. On the other hand, recent results from the adjacent Norwegian Skagerrak coast suggest that recruitment may vary between seasons as well as between years (Synnes *et al*., 2021; see also Jorde, Synnes *et al*., 2018).

Model selection suggests that the proportion of 0-group juveniles of the coastal ecotype is higher overall in Öresund than in the other ICES subdivisions, while decreasing with depth in all areas. The former inference might be expected, as biophysical modelling studies suggest that a large part of the juveniles assigned to the coastal ecotype may originate from the Öresund or southern Kattegat (Jonsson *et al*., 2016; Barth *et al*., 2017). That the coastal ecotype is more common in shallow areas is also expected, as bottom depth generally decreases toward the coast. However, the models with predictors including distance from the baseline did not perform as well as those including depth, suggesting a closer relationship of ecotype distribution with actual depth, rather than with distance from the baseline. For example, there were high proportions of offshore juveniles in both years at two stations within Gullmarsfjorden; Skår and Torgestad. While these two stations are located 13 and 21 km inshore from the baseline, they are substantially deeper than neighbouring stations, with bottom depths of 65 m and 107 m, respectively. The correlation with depth may thus be informative, and could reflect niche partitioning between the ecotypes in coastal areas. For example, niche partitioning related to depth (Grabowski *et al*., 2011; Michalsen *et al*., 2014) and differences in juvenile settling depths (Fevolden *et al*., 2012) have been observed between the migratory north-east Arctic cod (NEAC) and the stationary Norwegian coastal cod (NCC). The offshore-coastal divergences in beaked redfish, European seabass, and European anchovy are also associated with differences in depth distribution, as well as salinity tolerance (Allegrucci *et al*., 1997; Cadrin *et al*., 2010; Le Moan *et al*., 2016). The apparent utilisation of different depth strata may also be connected to differences in prey choice between the two cod ecotypes, which have been suggested for cod in Norwegian fjords (Kristensen *et al*., 2021).

Within the 2019 cohort, the coastal ecotype proportions increased from 0-group to 1-group in the inshore region. This might be the result of natural selection favouring the coastal ecotype in shallow, coastal environments, as suggested by Barth *et al.* (2019; see also “Environmental adaptation” below). On the other hand, the lack of 1-group cod in 2020 in several of the innermost fjord locations suggests that overall survival was low in inshore Skagerrak. Indeed, annual mortality rates as high as 75% have been suggested on the Norwegian Skagerrak coast (Olsen & Moland, 2011). However, the apparent “offshore shift” within the 2019 cohort could also have resulted from a net migration of 1-group cod toward the outer coastal zone. Different age classes of cod may well utilise different habitats (Fromentin *et al*., 2000; Pihl *et al*., 2006), but the habitat preference, feeding ecology, and behaviour of juvenile cod in this region are yet to be explored in detail.

### Environmental adaptation

The genetic variation at candidate loci suggests that the ecotypes may be genetically adapted to different environments. For instance, homozygotes for the valine allele (“homozygote 1” in Figure 5G) at the Hb-β1 locus are proposed to be more tolerant to hypoxic conditions, and prefer lower temperatures (Petersen & Steffensen, 2003; Brix *et al*., 2004). In addition, the migratory NEAC and the brackish-adapted eastern Baltic cod both have highly differentiated HbI genotypes compared to cod in other locations (Sick, 1961; Andersen *et al*., 2009), likely reflecting environmental adaptation. The higher frequency of the valine allele could thus provide the coastal ecotype with a higher tolerance for hypoxia and low temperatures. The ecotype differences in Hb-β1 allele frequencies observed in the present study were small, but consistent with a previous study comparing the Hb-β1 allele frequencies for cod in the North Sea and Kattegat (Andersen *et al*., 2009).

Similar to Hb-β1, the inversions on chr 2, 7, and 12 have also been associated with temperature and oxygen conditions, but also with salinity (Berg *et al*., 2015; Oomen, 2019). The ecotype-specific inversion state frequencies observed in this study were highly consistent with those from previous studies (Barth *et al*., 2017; 2019; Sodeland *et al.*, 2022), indicating that they are spatiotemporally stable in this geographical area. The most striking difference between ecotypes was in chr 12 inversion state, as indicated both by the large genotype frequency differences, and by most genome-wide outliers being located within this inversion. One of the chr 12 inversion states (“ancestral” in this study) was very rare in the coastal ecotype, a pattern highly consistent with previous studies (Barth *et al.*, 2017; 2019; Sodeland *et al*., 2022). Barth *et al.* (2019) showed that survival in the fjord environment is lower for homozygotes of the ancestral chr 12 state (“inverted” in Barth *et al*., 2019; see Table S3), which is more common within the offshore ecotype in the present study.

Model selection indicated that the inversion states on chr 2, 7, and 12 were not only different between ecotypes, but also correlated with ICES Subdivision, bottom depth, and/or distance from the baseline. This suggests that the coastal ecotype-like inversion genotypes are favoured in habitats that are shallower or located further inshore, the same areas where coastal ecotype juveniles were more frequent. Indeed, recent research suggest that the mode of natural selection (neutral, balancing, or directional) on the alternative inversion states partly depends on ecotype, but can also differ between locations (Sodeland *et al*., 2022). This motivates more research efforts aimed towards linking inversion genotypes to phenotypes and environmental variables. The inversions on chr 2, 7 and 12 have previously been broadly associated with salinity, temperature and oxygen conditions (Berg *et al.*, 2015; Kess *et al*., 2020). Experiments on cod larvae suggest that the inversions on chr 2 and 7 affect thermal plasticity and salinity tolerance, respectively, and that those on chr 2 and 12 are involved in cold-protection (Oomen, 2019). Hence, the associations with depth and/or distance from the baseline likely represent a more proximate relationship of the alternative inversion states to environmental differences between offshore and inshore, and deep and shallow, habitats. However, further studies may be required to assess the biological relevance of the low levels of genetic differentiation at the Hb-β1 and Hb-β5 loci, and the chr 2 and chr 7 inversions.

The outlier analysis provides additional evidence that differential environmental adaptation may underlie the offshore-coastal ecotype divergence. Two of the annotated outlier genes located within the chr 12 inversion, vitellogenin-2 (VTG2) and Tubulin-Tyrosine Ligase-Like Protein 7 (TTLL7), are in a genomic region characterised by a double crossover in Baltic cod (Matschiner *et al*., 2022). Both genes may be associated with environmental adaptation and reproduction, as VTG is involved in regulating fish egg buoyancy (Finn & Fyhn, 2010), and TTLL7 is involved in tubulin polyglutamylation, which may play a role in both sperm cells in cod testes (Klotz *et al*., 1999) and cold-adaptation of cod microtubules (Modig *et al.*, 1999). Vigilin (HDLP), which inhibits degradation of VTG mRNA (Cunningham *et al*., 2000), was also among the genes associated with outlier loci. Among the annotated outliers, there were also four genes with potential links to migratory behaviour (CACNB3-, OR2K2-, OR56A4-, and SLCO1C1-like). The calcium channel subunit CACNB3 is associated with the annotated outlier locus with the highest *F*_ST_ between the ecotypes, and has previously been linked with migratory phenotypes in fish (Cao *et al*., 2020; Gao *et al*., 2021). The two olfactory receptors OR2K2 and OR56A4 are located within a cluster of olfactory genes on chr 16, and olfaction is heavily involved in establishing long-term memory and homing behaviour in fish (Hara, 1994). SLCO1C1 is involved in thyroid hormone signalling (Admati *et al*., 2020), which may dictate the timing of spawning migrations in cod (Woodhead, 1959). SLCOC1 has previously been described as an outlier between “North Sea” and “fjord” cod in Norwegian Skagerrak (Barth *et al*., 2019). In addition, we detected outlier loci linked to genes associated with long-term memory (PPP1R14B; Cheng *et al*., 2015), social and acoustic behaviour (ISTR; Goodson & Bass, 2000), feeding behaviour and growth (PACAP; Xu & Volkoff, 2009, and LEP-R; Gorissen & Flik, 2014) in fish. Together, the functions of these genes provide mechanistic insights into how the divergence may have evolved and persisted between these sympatric ecotypes.

### Local coastal populations

There are at least three potential explanations for the dominance of the coastal ecotype in inshore areas. First, the fjords may receive more recruits originating from the known spawning grounds in the Kattegat and Öresund than from the North Sea (Stenseth *et al*., 2006; Jonsson *et al*., 2016; Barth *et al*., 2017). Second, the coastal ecotype may be better adapted to the environmental conditions associated with inshore habitats and therefore dominate these areas (Knutsen *et al*., 2018; Barth *et al*., 2019; Oomen, 2019). Third, the pattern might be attributed to local spawning populations inhabiting coastal areas that are closely related to Kattegat and Öresund cod (Svedäng *et al*., 2019).

In Norwegian fjords, local populations genetically similar to, but distinct from, cod from the Kattegat and Öresund have been identified (Barth *et al*., 2019), and both local spawning (Espeland *et al*., 2007; Jorde, Synnes *et al*., 2018) and highly resident behaviour (Knutsen *et al*., 2011; Kristensen *et al*., 2021) have been documented. Local spawning could potentially explain the dominance of coastal ecotype juveniles in the innermost portions of Swedish fjords (Färlevfjorden and Saltkällefjorden in particular, but also Åbyfjorden, Brofjorden, and Havstensfjorden). Recently, cod eggs of early developmental stages that assign genetically to adult fjord cod were documented also in Swedish fjords (Svedäng *et al*., 2019). As modelled drift velocities for eggs spawned in Öresund average < 5 km per day (Jonsson *et al*., 2016), the early (1-2 days old) egg stages found inside Swedish fjords likely originate from locally spawning adults (Svedäng *et al*., 2019). Moreover, models of pelagic egg drift on a local scale in Gullmarsfjorden and Brofjorden suggest that, if local spawning occurs, a high proportion of eggs are likely to be retained within fjords (P. Jonsson, pers. comm.; *cf*. Cianelli *et al*., 2010). Hence, the juveniles found inside fjords in the present study may originate from both locally spawning adults and adults spawning in areas outside of the fjords.

The analysis of 9956 genome-wide SNP loci, however, provided no evidence that coastal ecotype juveniles found inside Swedish fjords are genetically distinct from adult spawning populations in Kattegat and Öresund. Our sampling design included a high number of stations and a relatively low number of individuals per station, and this design was better suited to resolve the distribution of the ecotypes than to explore population structure within the coastal ecotype. Sequencing efforts with higher genomic resolution (such as whole-genome sequencing), more individuals per location, and more reference spawning adults would provide higher power to infer cryptic population structure. Nevertheless, if there are reproductively isolated local populations in the area, the level of genetic differentiation between them is likely very low and it may be restricted to specific genomic regions. In addition, even low migration rates that may have limited demographic importance in marine populations, can contribute with sufficient gene-flow to erode any genetic population structure (Allendorf *et al*., 2022). Hence, while the presence of genetic population structure is a strong indication of demographic independence, a lack thereof is not evidence of the opposite. Further studies assessing whether local spawning aggregations along the Swedish west coast represent demographically independent populations are therefore warranted.

### Conclusions and implications for management

Our results dispute the notion that cod along the Swedish Skagerrak coast generally originate from offshore spawning populations in the North Sea or Skagerrak (Svedäng, 2003; Cardinale & Svedäng, 2004). We show that there are considerable proportions of coastal ecotype juveniles, which should be accounted for in fisheries management, and in efforts to explain the underlying mechanisms of declining adult abundances. We suggest that future studies should look for alternative explanations connected to the population dynamics of the coastal ecotype, for instance whether adults perform spawning migrations to the Kattegat, Öresund or the Danish Belt Sea (Svedäng *et al*., 2007; André *et al*., 2016), or if larval and juvenile mortality has increased in the last decades. The lack of 1-group cod at the innermost fjord stations in this study coupled with declining adult abundances (ICES, 2021a-d) support the theory that shallow coastal habitats may have lost some of their function as a nursery for juvenile cod.

Altogether, this study provides an overview of the genetic population structure of Atlantic cod off the Swedish west coast. Our findings highlight the importance of treating the cod stock of the Skagerrak, Kattegat and Öresund as a mechanical mixture of two or more genetically distinct ecotypes. It is essential to consider this population structure and the local genetic adaptation for the conservation of Atlantic cod in this region. If genetic diversity is not restored in severely depleted cod stocks, it may negatively affect the potential for recovery of this ecologically and (once) economically important species.

## Supporting information

Supplementary material - Henriksson et al - bioRxiv

## Data availability statement

All data underlying the results presented in this study will be publicly available upon acceptance to a peer-reviewed journal.

## Acknowledgements

We thank Jakob Hemmer-Hansen for insightful comments on early versions of the manuscript. Sampling and genetic analysis were supported by contract Dnr 1639-2020 within 1:11 Åtgärder för havs-och vattenmiljö from SwAM. Sequencing was performed by the SNP&SEQ Technology Platform in Uppsala. The facility is part of the National Genomics Infrastructure (NGI) Sweden and Science for Life Laboratory. The SNP&SEQ Platform is also supported by the Swedish Research Council and the Knut and Alice Wallenberg Foundation. Further funding was provided by the EU Interreg project MarGen II, and the study was performed within the Linnaeus Centre for Marine Evolutionary Biology.

