## Supplementary material - Henriksson et al - bioRxiv for "Mixed origin of juvenile Atlantic cod (*Gadus morhua*) along the Swedish west coast": Supplementary notes and figures - Henriksson et al - bioRxiv.pdf

#### S1. Quality control and filtering

Loci that were unsuccessfully genotyped (n = 7) or monomorphic (n = 4) for this set of individuals were filtered out. To control for genotyping errors, we then tested all loci in the diagnostic panel for deviations from Hardy-Weinberg (HW) proportions, using the R package PEGAS (v0.14; Paradis, 2010). To avoid filtering out informative loci, we first assigned individuals to ecotypes (see "Ecotype assignment") and only removed loci deviating significantly from HW proportions within both ecotypes (n = 2). All locus pairs in the ecotype-diagnostic panel were also analysed for linkage disequilibrium (LD), using the DARTR package (v1.9.4; Gruber et al., 2018), but no pair of loci exceeded our threshold at $R^2 > 0.1$ in both ecotypes. Individuals genotyped successfully for less than 80% of the final ecotype-diagnostic set of loci (< 27 loci) were removed from downstream analysis (n = 15), along with one reference individual from Gullmarsfjorden with conflicting genotyping compared to a previous study.

#### S2. Evaluation of ecotype assignment

The ecotype assignment using K-means clustering was evaluated using four different methods. First, we performed discriminant analysis of principal components (DAPC) in ADEGENET (v2.1.3; Jombart, 2008) to calculate posterior probabilities of the assignment. Second, we performed a principal component analysis (PCA) on both the filtered ecotype-diagnostic SNP panel and a set of genome-wide loci (see "Genome-wide loci") with the ADE4 R package (v1.7-16; Dray & Defour, 2007) to visualise between-group differences. Third, we calculated ancestry coefficients using sparse non-negative matrix factorization (sNMF) for K = 1 – 10 and 50 repetitions for each K value, using the LEA package (v3.1.2; Frichot & François, 2015). The optimal K was selected as the K causing the sharpest decrease in cross-entropy compared to K-1, similar to above. Fourth, we used the ASSIGNPOP package (v1.2.2; Chen *et al.*, 2020), with default settings, to calculate assignment probabilities for a "North Sea" or "Kattegat/Öresund" origin by treating adult cod from these areas as reference populations.

Both sequential K-means clustering in ADEGENET, and sNMF indicated an optimal K = 2 (Figures S4-S5), supporting our assumption of two groups (the offshore and coastal ecotype) present in the dataset. The assigned ecotype corresponded well with posterior probabilities from the DAPC, as 99.2 (487/491) and 99.4 % (493/496) of individuals assigned to the offshore and coastal ecotypes, respectively, had posterior probabilities above 50 % for their assigned ecotype (Figure S3A). In the sNMF analysis, 99.0 (486/491) and 95.8 % (475/496) of offshore and coastal individuals, respectively, had an ancestry coefficient above 50 % for their assigned ecotype (Figure S3B). In ASSIGNPOP, similarly, 90.8 % (426/469) and 99.6 % (455/457) of the offshore and coastal juveniles, respectively, had assignment probabilities above 50 % for the adult population corresponding to the assigned ecotype (Figure S3C). We used K-means clustering as the main ecotype assignment method as this method makes less assumptions regarding the behaviour of the genetic markers used, and is not dependent on reference populations with relatively small sample sizes, compared to the alternative assignment methods.

#### S3. Model selection

The distance from the baseline was calculated in ArcMap™ (v.10.7.1; ArcGIS®, Esri) by creating a barrier-aware Euclidean distance raster from the Swedish (https://www.sjofartsverket.se/sv/tjanster/sjokortsprodukter/kopa-sjokort2/sjokort/havsgranser/baslinjer/), Norwegian, and Danish baselines (Flanders Marine Institute, 2019) with the coast as a barrier layer. The cell size used to compute the distance raster was $8 \times 10^{-4}$

decimal degrees for offshore stations and $4 \times 10^{-4}$ decimal degrees for inshore coastal stations, to account for the complexity of the coastline. Station-wise distances were obtained by extracting point values from the distance raster using the station coordinates. The distances were transformed into negative values for stations located offshore of the baseline, and positive for stations located inshore of the baseline.

To avoid overfitting and to minimise the associated risk of spurious findings, we plotted all potential two-way interactions and only included an interaction in the pool of explanatory variables if an effect on the dependent variable was indicated by these plots. We then fitted regression models with all possible combinations of the included explanatory variables as predictors. To minimise multicollinearity within predictors, we removed all fitted models that included explanatory variables with a variance inflation factor (VIF) > 2, using the CAR package (v3.0-12; Fox & Weisberg, 2019). To enable calculating VIF for the ordinal regression models, they were temporarily reformulated as linear regression models for this filtration step. Further, the linear regression models on ecotype and sex were filtered out if they did not meet the assumption of homoscedasticity, according to a Breusch-Pagan test in the LMTEST package (v0.9-38; Zeileis & Hothorn, 2002). We also controlled that the 10 highest-ranked linear models were insensitive to influential datapoints and that residuals were normally distributed, using the STATS package. After the multicollinearity filtering, the ordinal regression models were filtered out if they did not meet the assumption of proportional odds, according to a Brant test as applied in the BRANT package (v0.3-0; Schlegel & Steenbergen, 2020).

### S4. Genotyping and assignment

The use of genetic markers with above-average genetic differentiation offers a powerful and cost-efficient method to distinguish populations (Nielsen *et al.*, 2012). However, these high-graded markers are often linked to genes that may be under selection, which brings uncertainty to whether the inferred genetic structure using these markers represents demographic population structure or is merely an effect of selective mortality (Funk *et al.*, 2012; Gagnaire *et al.*, 2015; Hellberg, 2009). Moreover, the ecotype-diagnostic SNP panel used in the present study was developed specifically to identify the two ecotypes, so the risk of ascertainment bias is inherently high; that is, any potential third population or ecotype would not be identified by these markers. In contrast, the genome-wide loci obtained through 2b-RAD sequencing consist of polymorphic SNPs randomly distributed across the genome. The consistency of our ecotype assignment with the PCA clusters using the genome-wide markers thus provides an independent validation that our ecotype-diagnostic SNP panel accurately describes the demographic population structure, and that only two major clusters were present. The PCA clusters also persisted when loci under putative selection were removed, which corroborates previous studies indicating a low, yet temporally stable, neutral genetic differentiation between the two ecotypes (Knutsen *et al.*, 2011). In addition, the individual assignment to ecotype was highly consistent between different assignment methods, including K-means clustering, ancestry analysis, and population assignment (Figure S3; Table S6). Hence, this study supports the conclusions of Jorde *et al.* (2018), that high-graded SNP panels such as the one used here can be trusted in inferring demographic population structure of Atlantic cod in this area.

### S5. Inversion scans

The inversion scans with the INVERSION package (v1.40.0, Cáceres, 2021) were done by coding haplotype blocks of 3 SNPs flanking each candidate breakpoint, and then scanning each chromosome for inversions with window sizes of 1-30 Mb, setting a maximal number of 30 iterations for the expectation-maximisation algorithm. The minimum minor allele frequency allowed was 5 % and all models with a BIC > 0 were evaluated. In cases where the listed breakpoint values differed for the returned inversion models, we defined the most extreme values as our inversion breakpoints (Table S17). This conservative approach was taken to minimise the risk of including SNP loci linked to inversions in subsequent analyses.

The inversion scans detected the previously described inversions on chr 2, 7, and 12, and the breakpoints coincided with large genomic blocks with elevated LD (Figure S6). The inverted region on chr 1 was not identified by these scans, likely because all individuals were of the same chr 1 inversion state, according to the targeted SNP genotyping. The inversion scans indicated that 538 SNP loci (119 on chr 2, 217 on chr 7, and 202 on chr 12) were located inside of inverted regions, in total.

The reduced panel of SNPs used to assign inversion state also appeared to have a high accuracy for most individuals, as the assigned inversion states were consistent with those inferred using the genome-wide loci (Figure S7; Table S6). However, the few individuals with conflicting genotyping using the targeted panel (one homozygous locus and one heterozygous locus within the same inversion; “Mixed” in Figure S2, S7, and Table S18) did not appear to have intermediate inversion states, but were all either homo- or heterozygous for the inversions according to the genome-wide markers. This suggests that future panels would benefit from including a higher number of SNP loci per inverted region, to increase the power to resolve potential genotypic inconsistencies.

### References, supplementary notes


Cáceres, A. (2021). *inveRsion: Inversions in genotype data. R package version 1.40.0*. https://doi.org/doi:10.18129/B9.bioc.inveRsion

Chen, K.-Y., Marschall, E. A., Sovic, M. G., Fries, A. C., Gibbs, H. L., & Ludsin, S. A. (2020). *assignPOP: Population Assignment using Genetic, Non-Genetic or Integrated Data in a Machine Learning Framework*. *R package version 1.2.2*. https://CRAN.R-project.org/package=assignPOP

Dray, S., & Dufour, A. (2007). The ade4 Package: Implementing the Duality Diagram for Ecologists. *Journal of Statistical Software, 22*(4), pp. 1-20. https://doi.org/10.18637/jss.v022.i04

Flanders Marine Institute (2019). *Maritime Boundaries Geodatabase: Internal Waters, version 3*. https://doi.org/10.14284/385

Fox, J. & Weisberg, S. (2019). *An {R} Companion to Applied Regression,* Third Edition. Thousand Oaks CA: Sage. URL: https://socialsciences.mcmaster.ca/jfox/Books/Companion/

Frichot, E., & François, O. (2015). LEA: An R package for landscape and ecological association studies. *Methods in Ecology and Evolution, 6*(8), 925-929. https://doi.org/10.1111/2041-210X.12382

Funk, W. C., McKay, J. K., Hohenlohe, P. A., & Allendorf, F. W. (2012). Harnessing genomics for delineating conservation units. *Trends in ecology & evolution, 27*(9), 489-496. https://doi.org/10.1016/j.tree.2012.05.012

Gagnaire, P. A., Broquet, T., Aurelle, D., Viard, F., Souissi, A., Bonhomme, F., Arnaud-Haond, S., & Bierne, N. (2015). Using neutral, selected, and hitchhiker loci to assess connectivity of marine populations in the genomic era. *Evolutionary applications, 8*(8), 769-786. https://doi.org/10.1111/eva.12288

Gruber, B., Unmack, P. J., Berry, O. F., & Georges, A. (2018). dartr: An r package to facilitate analysis of SNP data generated from reduced representation genome sequencing. *Molecular Ecology Resources, 18*(3), 691-699. https://doi.org/10.1111/1755-0998.12745

Hellberg, M. E. (2009). Gene flow and isolation among populations of marine animals. *Annual Review of Ecology, Evolution, and Systematics, 40*, 291-310. https://doi.org/10.1146/annurev.ecolsys.110308.120223

Jombart T. (2008) adegenet: a R package for the multivariate analysis of genetic markers *Bioinformatics 24,* 1403-1405. https://doi.org/10.1093/bioinformatics/btn129

Jorde, P. E., Synnes, A. E., Espeland, S. H., Sodeland, M., & Knutsen, H. (2018). Can we rely on selected genetic markers for population identification? Evidence from coastal Atlantic cod. *Ecology and Evolution, 8*(24), 12547-12558. https://doi.org/10.1002/ece3.4648

Knutsen, H., Olsen, E. M., Jorde, P. E., Espeland, S. H., André, C., & Stenseth, N. C. (2011). Are low but statistically significant levels of genetic differentiation in marine fishes ‘biologically meaningful’? A case study of coastal Atlantic cod. *Molecular ecology, 20*(4), 768-783. https://doi.org/10.1111/j.1365-294X.2010.04979.x

Nielsen, E. E., Cariani, A., Mac Aoidh, E., Maes, G. E., Milano, I., Ogden, R., Taylor, M., Hemmer-Hansen, J., Babbucci, M., Bargelloni, L., Bekkevold, D., Diopere, E., Grenfell, L., Helyar, S., Limborg, M. T., Martinsohn, J. T., McEwing, R.,

Panitz, F., Patarnello, T., ... & Carvalho, G. R. (2012). Gene-associated markers provide tools for tackling illegal fishing and false eco-certification. *Nature communications*, *3*(1), 1-7. https://doi.org/10.1038/ncomms1845

Paradis, E. (2010). pegas: an R package for population genetics with an integrated–modular approach. *Bioinformatics, 26*(3), 419-420. https://doi.org/10.1093/bioinformatics/btp696

https://www.sjofartsverket.se/sv/tjanster/sjokortsprodukter/kopa-sjokort2/sjokort/havsgranser/baslinjer/

Schlegel, B. & Steenbergen, M. (2020). *brant: Test for Parallel Regression Assumption. R package version 0.3-0*. https://CRAN.R-project.org/package=brant

Zeileis, A., & Hothorn, T. (2002). Diagnostic Checking in Regression Relationships. *R News 2*(3), 7-10. URL https://CRAN.R-project.org/doc/Rnews/

### Supplementary figures

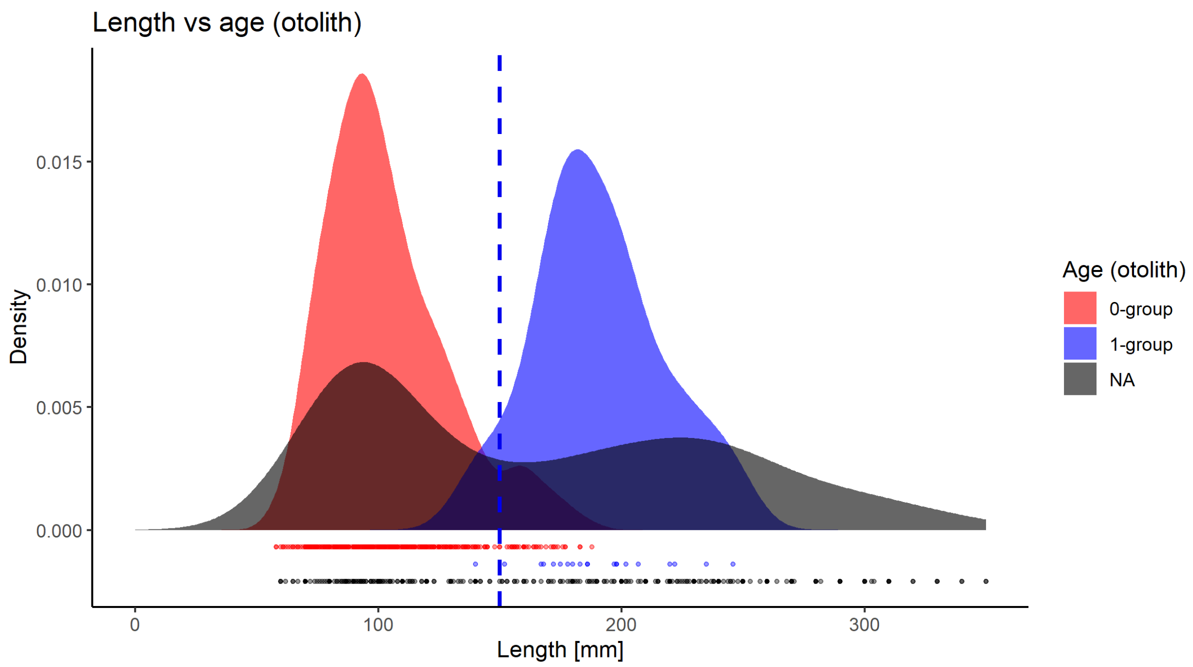


**Figure S1.** Length distribution of the juveniles included in the study. Colour corresponds to the age, as determined through otolith readings, and individuals that have not been aged are labelled “NA”. The horizontal dashed line indicates the cut-off value of 150 mm, used for cohort assignment of juveniles that had not been aged based on otolith readings.

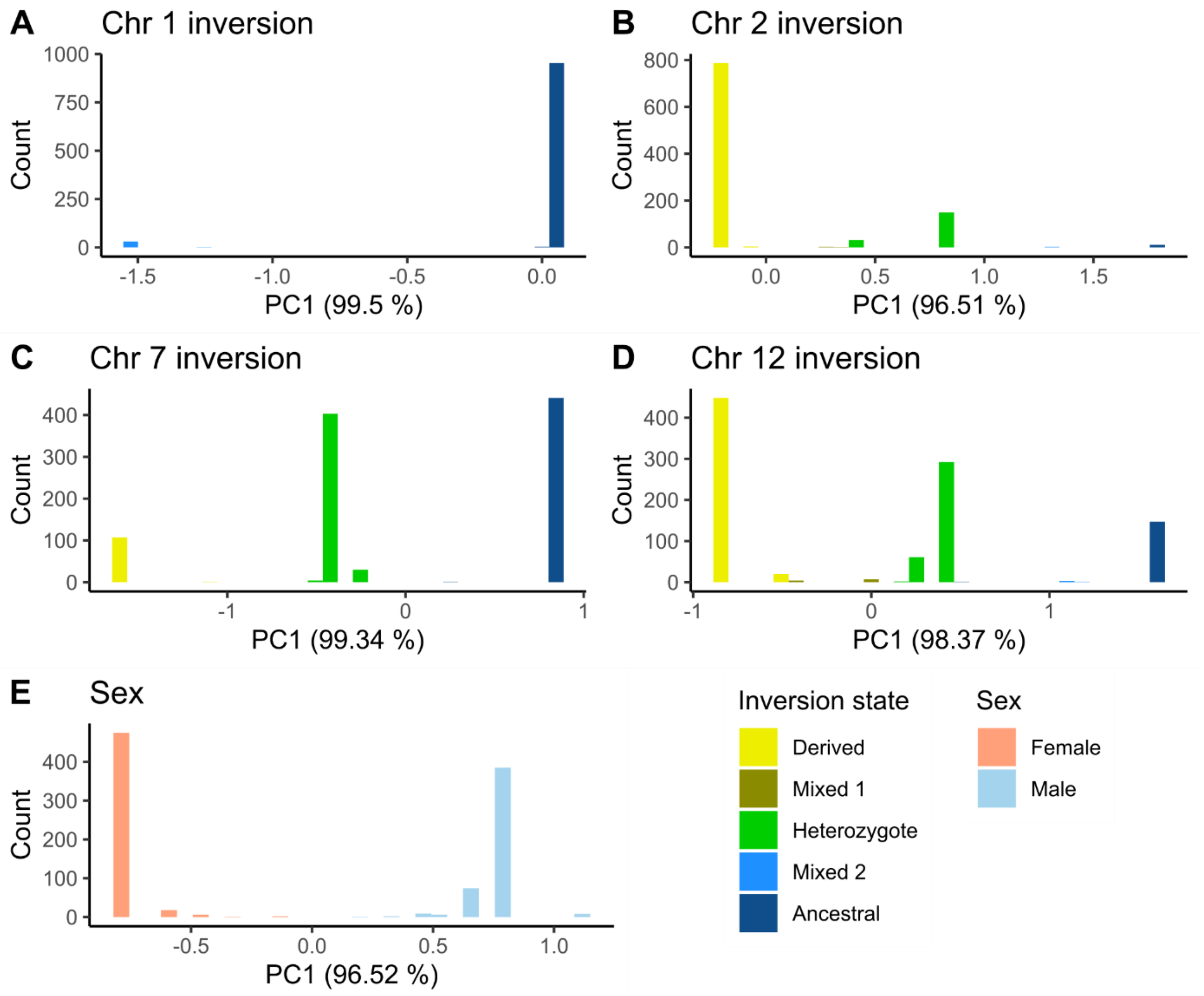


**Figure S2.** Histograms of individual PC1 coordinates upon which the assignment of **A-D)** inversion state and **E)** sex was based. The PCAs were performed using loci from the targeted SNP panel. Individuals with one homozygous and one heterozygous locus within the same inversion are labelled “Mixed” and were excluded from downstream analyses.

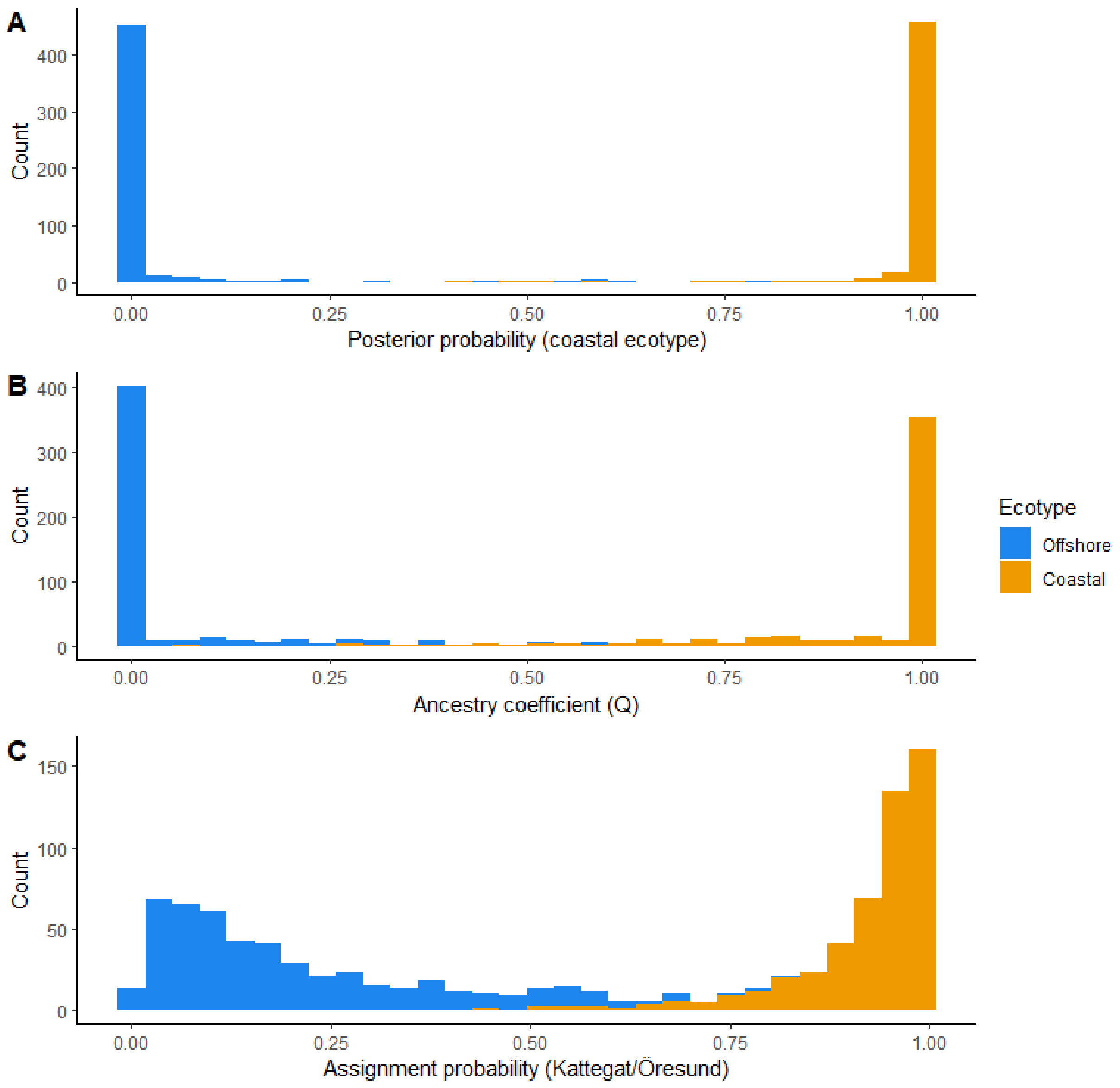


**Figure S3.** Histograms of **A)** posterior group assignment probabilities in DAPC, **B)** ancestry coefficients from the sNMF analysis, and **C)** assignment probabilities for the reporting group including Kattegat/Öresund adults in ASSIGNPOP. Colour corresponds to the assigned ecotype according to the K-means clustering.

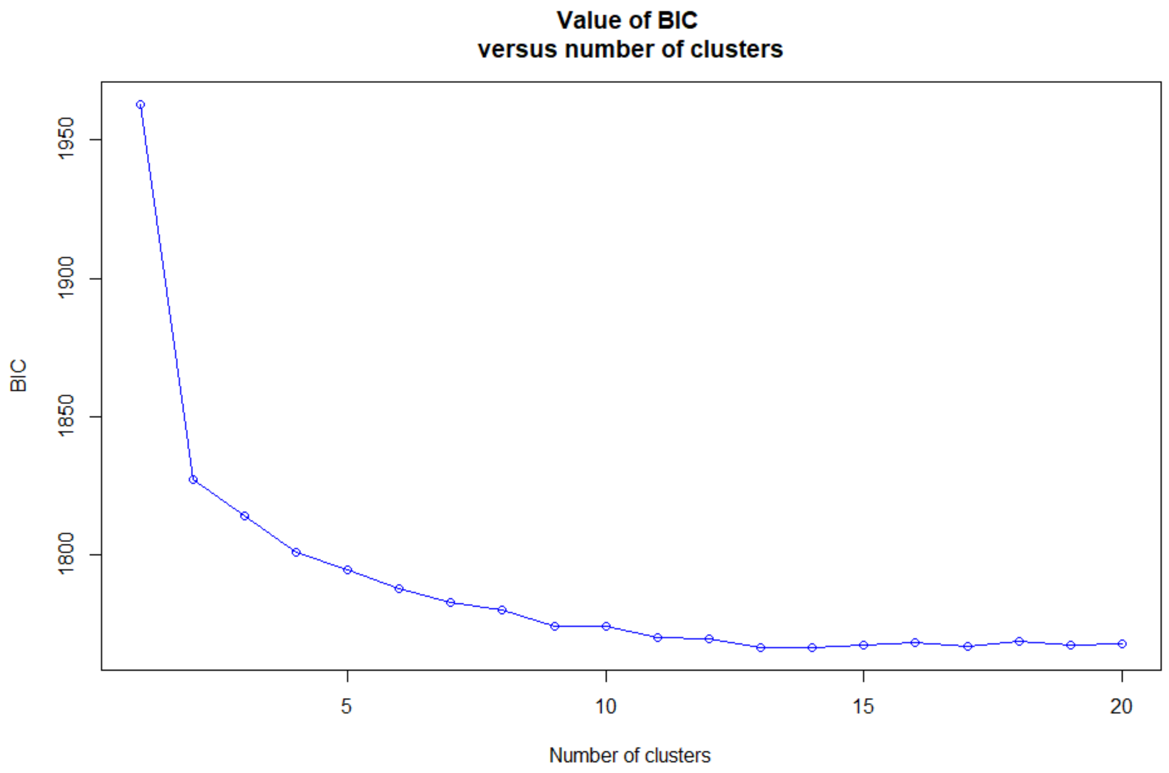


**Figure S4.** Sequential K-means clustering based on genotypes at the 33 ecotype-diagnostic loci. Bayesian Information Criterion (BIC) values shown for K = 1-20.

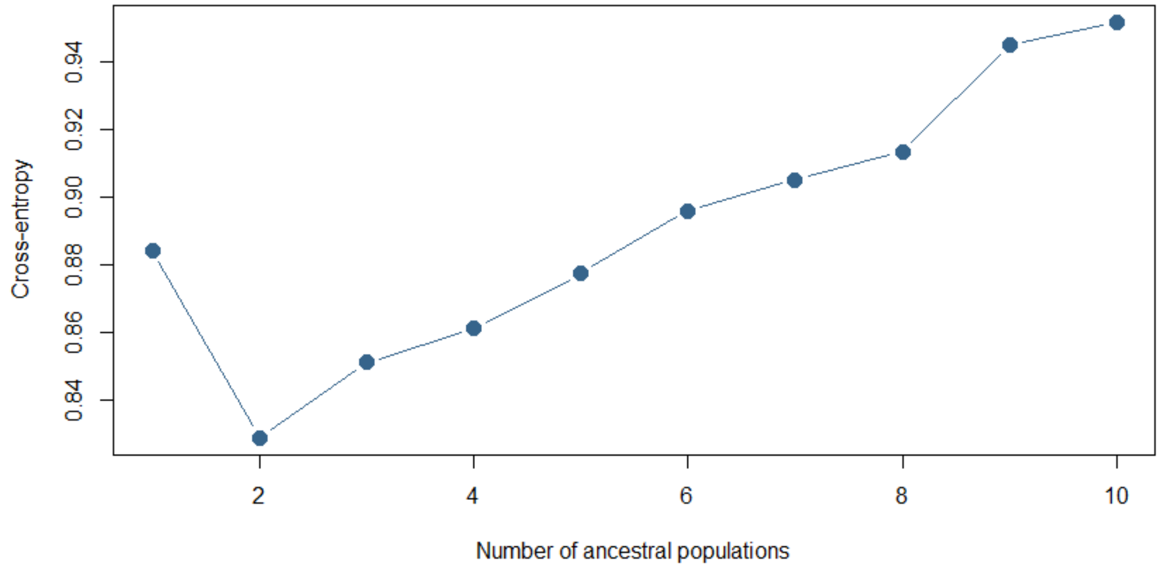


**Figure S5.** Results of the sparse non-negative matrix factorisation (sNMF) based on the 33 ecotype-diagnostic loci. Cross-entropy values shown for K = 1-10.

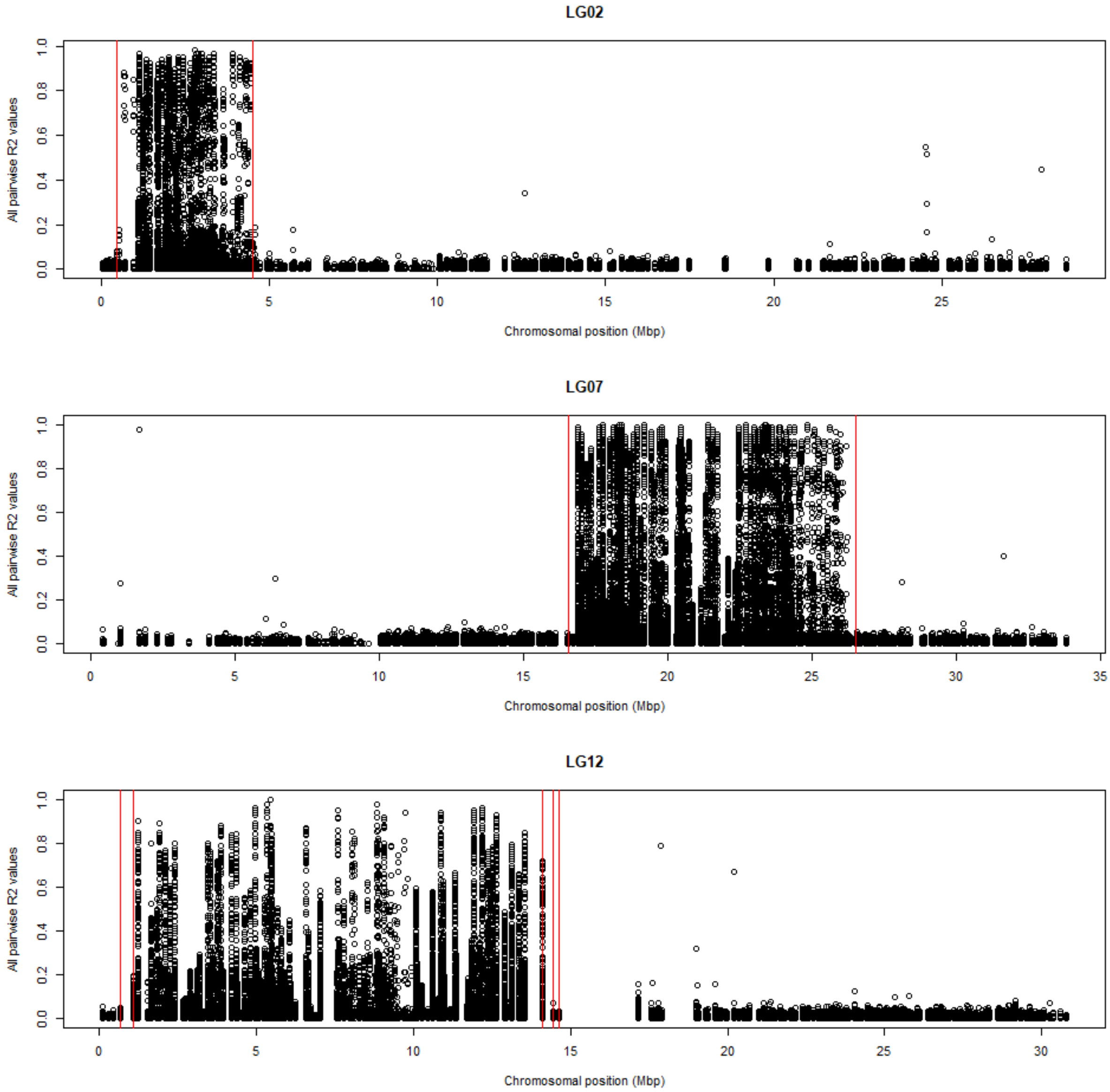


**Figure S6.** Pairwise $R^2$ values indicating the linkage disequilibrium of genome-wide SNP loci on LG2,7, and 12. The red lines indicate the most extreme inversion breakpoints for inversion scans with window sizes of 1-30 Mbp, using the inveRsion package. For LG2 and 7, the same inversion breakpoints were returned regardless of window size. For LG12, a total of five different inversion breakpoints were returned. We used the most extreme (left-most and right-most) breakpoints for our downstream analyses.

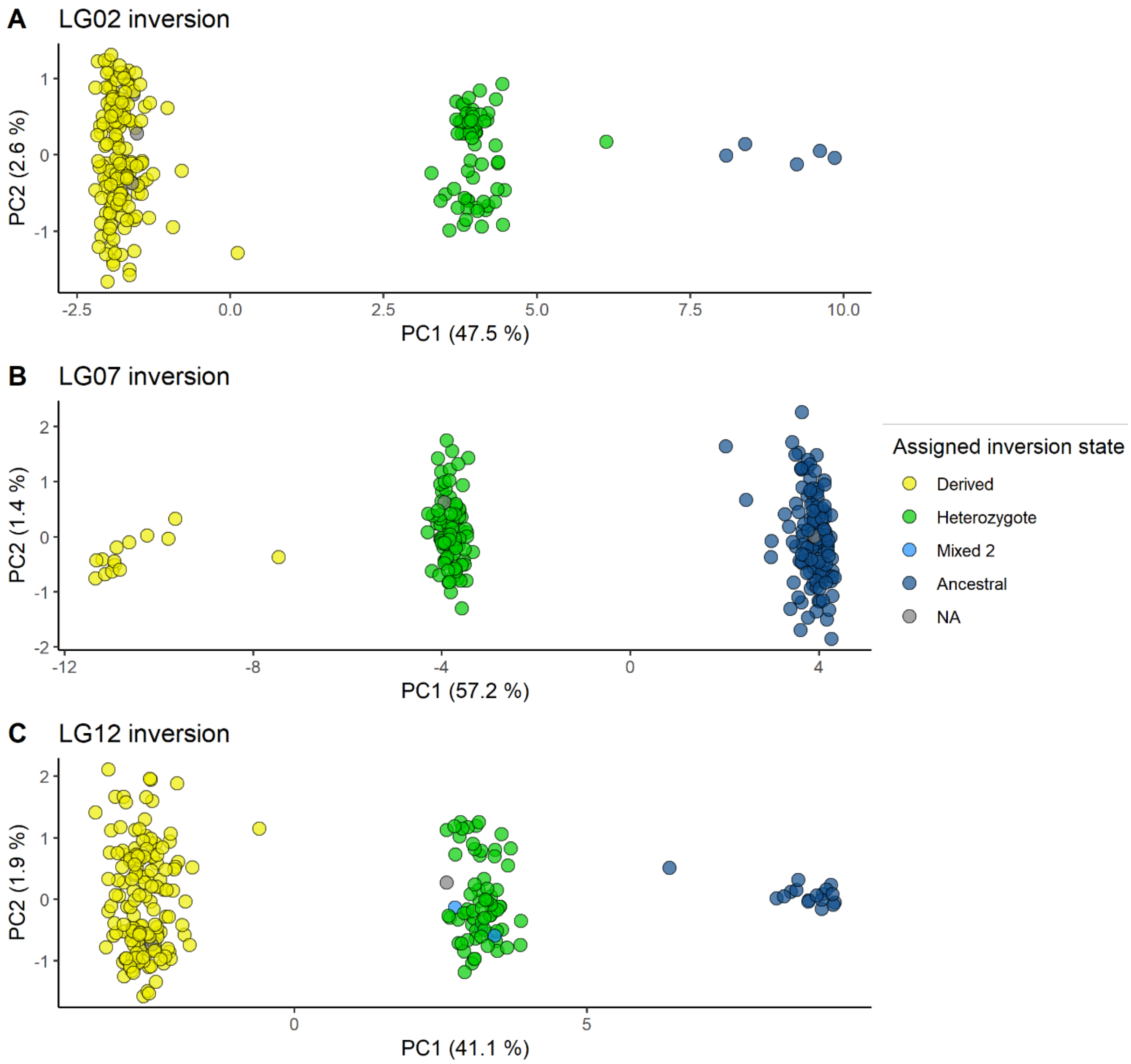


**Figure S7**. PCA clustering of genome-wide loci located within inversions, with colour indicating the assigned inversion state using the inversion-diagnostic SNPs from the panel of targeted SNPs. Individuals with one homozygous ancestral and one heterozygous locus within the same inversion are labelled "Mixed 2". Three individuals who were excluded from ecotype assignment due to low genotyping success for the ecotype-diagnostic loci, but that are included in this figure, are labelled "NA".
